# Locat: Joint enrichment and depletion testing identifies localized marker genes in single-cell transcriptomics

**DOI:** 10.64898/2026.04.03.716370

**Authors:** Wesley Lewis, Yariv Aizenbud, Francesco Strino, Yuval Kluger, Fabio Parisi

## Abstract

Several methods identify marker genes that delineate cell populations in single-cell transcriptomic data, yet most emphasize enrichment within candidate populations without testing whether expression is significantly reduced elsewhere. We present Locat, a framework for identifying highly specific localized genes by testing whether expression is concentrated within compact regions of a cellular embedding and depleted outside them. For each gene, Locat fits weighted Gaussian mixture models to gene-specific and background densities, computes concentration and depletion statistics, and integrates them into a unified localization score. Across synthetic benchmarks with controlled ground truth, Locat detects uni-modal, multi-modal, and sparse localized patterns and loses significance when expression becomes indistinguishable from background structure. In developmental, perturbation, and differentiation datasets, Locat identifies compact marker sets that capture lineage organization, condition-specific programs, and temporal dynamics. These sets are often smaller than highly variable gene selections, while embeddings built from them preserve major cell populations and developmental programs in several cases. In murine dermis, interferon-treated PBMCs, and retinoic acid-induced embryonic stem cell differentiation, localized genes recover differentiation trajectories, stimulus-responsive programs, and reproducible stage-specific patterns. Together, these results show that jointly assessing concentration and depletion yields specific, interpretable marker genes.

## 1 Introduction

A central goal in single-cell transcriptomics is to identify marker genes whose expression serves as a reliable indicator of biologically meaningful cell populations and states. These markers serve multiple purposes. They can distinguish discrete cell types such as T cells and B cells, trace continuous developmental trajectories such as hematopoietic differentiation, reveal functional heterogeneity within apparently uniform populations such as activated and quiescent states, and define specialized subsets within major lineages, such as naive, regulatory, and memory CD4+ T cells. In many cases, these subsets occupy contiguous regions in transcriptional space despite sharing broader lineage identity.

In this context, localized marker genes exhibit two defining properties: expression that is enriched within a compact subset of cells, together with marked depletion elsewhere. This depletion, particularly in substantial regions outside the target population, is essential for biological specificity and reliable cell type identification. Consistent with this view, a recent large-scale benchmarking study evaluating marker genes across diverse biological contexts shows that marker utility depends critically on expression specificity relative to the full cellular background [1]. In particular, genes with expression confined to restricted regions or cell states, and little to no expression elsewhere, are favored over broadly or diffusely expressed genes that exhibit local enrichment [1]. In practice, most marker discovery methods prioritize genes based on enrichment criteria, favoring expression patterns that are concentrated, smooth, or compact on a cell–cell graph, or enriched within clusters of similar cells. These approaches typically assess enrichment without explicitly testing for the absence of expression outside candidate regions. As a result, diffuse expression patterns cannot be readily distinguished from truly localized ones. Consequently, genes with high baseline expression may rank highly under enrichment-based scoring schemes despite lacking a well-defined localized domain, conflating statistical enrichment with biological specificity. These limitations motivate the need for approaches that jointly assess enrichment within candidate regions and exclusion outside them, enabling the identification of genes that demarcate cellular populations rather than merely correlate with them.

We present Locat, a statistical framework that formalizes this dual requirement by testing for both concentrated enrichment (which we term *concentration*) and significant exclusion from the remaining cellular background (which we term *depletion*). Rather than relying on predefined cell groupings or discrete partitions, Locat models gene expression patterns as continuous probability densities within user-supplied embeddings, such as PCA or Uniform Manifold Approximation and Projection (UMAP) representations. For each gene *g*, Locat fits a gene-specific weighted Gaussian mixture model (WGMM) to estimate the density of cells expressing that gene and contrasts this with a background WGMM describing the overall distribution of all cells in the embedding. Concentration and depletion are then assessed by comparing these two density models: the concentration test quantifies the extent to which the gene-specific density aggregates into compact regions of the embedding, while the depletion test evaluates whether expression is significantly under-represented across a substantial amount of the background density outside these concentrated regions. The concentration and depletion *p*-values are first combined using Cauchy aggregation. This Cauchy-combined value is then adjusted to yield the final localization score used for thresholding and cross-dataset comparisons. By construction, the adjusted score is conservative relative to the Cauchy-combined value (see Methods). Because gene-level tests share a common embedding and background model and differ substantially in prevalence, we assess calibration empirically using simulation-based null analyses across the range of prevalences encountered in the data. We therefore do not rely on a single genome-wide correction as the primary calibration device (Supplementary Methods S1.3; Supplementary Fig. S1).

Conceptually, Locat belongs to a broad class of marker discovery methods that seek genes whose expression is concentrated within restricted regions of transcriptional space, whether those regions are defined by clusters, cell–cell graphs, or continuous manifolds. Several existing tools formalize this idea using different statistical criteria. singleCellHaystack identifies genes with non-uniform expression distributions by comparing observed and expected densities using Kullback–Leibler divergence [2]. Hotspot identifies genes whose expression exhibits significant local autocorrelation on a cell–cell similarity graph, highlighting genes whose values are predictable from their transcriptional neighborhood rather than randomly distributed [3]. GiniClust uses inequality metrics to prioritize genes enriched in rare populations [4]. Graph-diffusion–based methods such as GSPA and LMD operate directly on cell–cell manifolds to capture smooth, coherent, or compact expression patterns [5, 6]. While these approaches differ in how regions are represented and scored, they are unified by an emphasis on enrichment or structured variability as the primary criterion for marker selection.

We apply Locat across diverse biological systems, including embryonic dermis differentiation, perturbation experiments, and temporal developmental series. Across these settings, the dual assessment of enrichment and depletion identifies highly specific markers that capture both discrete cell types and continuous biological processes, reveal condition-specific localization patterns, and track the temporal emergence of localized expression during development. In embryonic differentiation and perturbation data, embeddings constructed using only top localized genes recover the major cell types and continuous trajectories observed in highly-variable gene embeddings, while relying on a smaller and more interpretable set of markers. In comparisons with existing clustering-free marker selection methods, Locat identifies gene sets that are notably distinct from those prioritized by enrichment-only approaches, with discrepancies driven by broadly expressed background genes filtered by Locat and by selective markers not emphasized by enrichment-based criteria. In time-series data, Locat detects genes whose localization emerges dynamically over time, and localization patterns are consistent across replicates at the same time point, supporting comparisons of localized programs across samples and conditions.

Together, these results show that Locat provides a general and reliable framework for discovering highly specific localized markers. By directly testing both enrichment within a region and depletion outside it, Locat produces interpretable, high-specificity gene sets that support meaningful comparisons across samples, conditions, and datasets.

## 2 Results

### 2.1 Localization requires selective absence

In marker gene discovery, scRNA-seq data is typically represented in a chosen representation of cells, such as tissue coordinates, a low-dimensional latent space (PCA, tSNE or UMAP), or a cell–cell neighborhood graph. In any of these representations, a gene can be locally enriched in this space without being a reliable marker. The key point of failure for existing tools is background expression: a gene may peak in one region yet remain detectably expressed across all other regions. Such a gene correlates with structure but does not identify or demarcate a particular cell population.

This motivates a stricter operational definition of localization. A localized marker should exhibit two complementary properties: (i) concentrated expression in compact regions of high density (concentration), and (ii) selective absence from substantial portions of the remaining cell population, particularly where many cells reside (depletion). The schematic in Fig. 1 illustrates why these properties are not equivalent: “enriched” describes a local maximum, whereas “enriched and depleted” describes a maximum paired with meaningful contrast against the global background density. Only the latter provides the necessary specificity required for population demarcation.

**Fig. 1.**
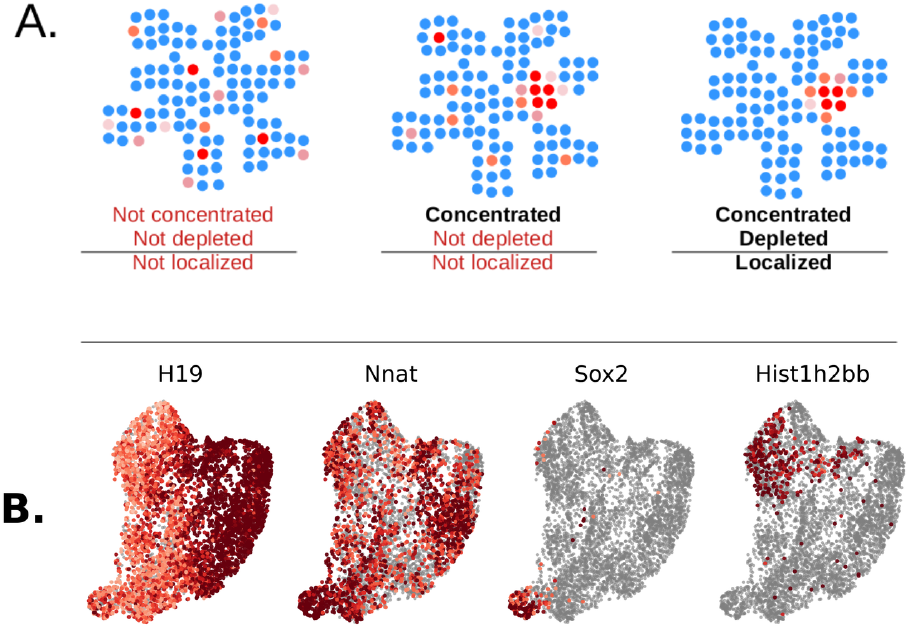
Illustrative examples of localized and not localized gene expression profiles. **A**. Three schematic genes on the same manifold (each dot a cell, red expressing and blue non-expressing). The left gene is not concentrated. The middle gene is concentrated but not depleted outside its enriched region. The right gene is both concentrated and depleted, which defines localization. **B**. UMAP embeddings from a murine dermal condensate dataset illustrating concentration without localization (*H19, Nnat*) and concentration with depletion (*Sox2, Hist1h2bb*).

We illustrate this distinction using murine embryonic dermis data from [7] and four representative genes in Fig. 1. These examples are chosen solely to demonstrate the conceptual cases below, whereas a more exhaustive evaluation of the dataset is given in Sections 2.3, 2.5, and 2.6.

- Concentrated only: *H19, Nnat*. Expression is clearly concentrated into a compact region, but appreciable expression remains visible across other populated regions of the embedding. These genes look localized under enrichment-first scoring because they have a strong peak, yet they do not cleanly define a population boundary.
- Concentrated and depleted: *Sox2, Hist1h2bb*. Expression is concentrated into a compact domain and is also markedly absent elsewhere, including in regions where many cells reside. Compared to *H19* and *Nnat*, these genes are better markers in the practical sense: presence is informative and absence is informative.

Depletion is thus not an incremental modification of enrichment scoring, but a necessary criterion for marker specificity. Having established the conceptual distinction between concentration and depletion, we formalize both as hypothesis tests for each individual gene in a fixed low-dimensional cell embedding, where proximity reflects similarity in transcriptional state. The concentration test evaluates whether a gene’s expression mass is more aggregated within this embedding than expected under a null model in which expressing cells are randomly distributed according to the background transcriptional activity.

#### Concentration

We quantify concentration by contrasting two density functions over the embedding. The first, *f*_*g*_, represents the density of gene *g*’s expression, and the second, *f*_0_, represents the background cell density. We define the concentration score as:

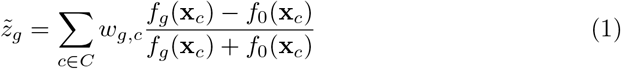

where *C* denotes the set of all cells and 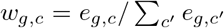 denotes the normalized expression weight of gene *g* in cell *c* (Section 4). The symmetric difference form (*f*_*g*_ − *f*_0_)*/*(*f*_*g*_ + *f*_0_) is bounded in [−1, 1] and measures the relative deviation of gene density from background at each cell location. The score aggregates these deviations across all expressing cells: positive values indicate that expression concentrates in regions of elevated density relative to the null expectation.

We estimate *f*_*g*_(**x**) from cells expressing gene *g*, weighted by that gene’s expression, and we estimate *f*_0_(**x**) from all cells in the embedding (see Section 4.1.2). The full operational definition of the concentration statistic is given in Section 4.1.3, including prevalence weighting and normalization. Significance is then obtained by calibrating this statistic against prevalence-matched randomized expression patterns, yielding a concentration *p*-value that controls for expression prevalence.

#### Depletion

We quantify depletion by testing whether gene expression is under-represented across a substantial portion of the embedding density. For each gene *g* and contrast threshold *λ* ≥ 1, we define a region of depletion

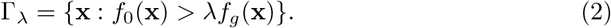

Within this region of depletion, we compute the observed expressing-cell fraction and the background density fraction

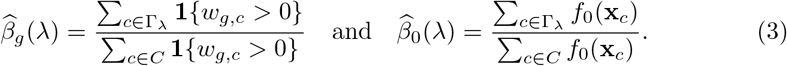

Intuitively 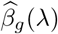 measures the fraction of gene *g*’s expressing cells that lie in regions where background density substantially exceeds gene density, while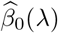 measures the fraction of overall cellular density in those same regions.

Depletion is assessed by testing whether

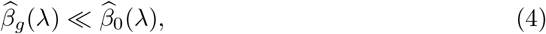

i.e., whether the gene avoids regions where many cells reside. The resulting depletion statistic corresponds to the tail probability under a null model in which gene expression follows the background density (see Section 4.1.4).

In practice, we evaluate depletion for different values of *λ* ∈ Λ (see Section 4.1.4) to capture depletion at different stringency levels, apply effective sample size corrections to account for autocorrelation of neighboring cells, and adjust for multiple testing across thresholds. The final depletion *p*-value represents the minimum across thresholds after correction.

#### Joint Testing: Localization

Locat evaluates gene localization by jointly testing for concentration and depletion (see Section 4.1). A gene is considered localized only if it shows significant evidence for both aggregation/compactness and exclusion from large portions of transcriptional space (see Section 4.1.5), aligning the statistical criterion with the biological goal of acting as a compact population indicator.

To motivate clear examples of localized genes, we again consider four genes from murine embryonic dermal condensates dataset [7]. Under our criterion, *Sox2* and *Hist1h2bb* are prioritized because they satisfy both requirements, whereas *H19* and *Nnat* fail the selective-absence criterion despite exhibiting strong local enrichment. All four genes show highly significant concentration (*p <* 10^*−*6^).

For *Sox2* and *Hist1h2bb*, depletion is strongly supported (depletion *p <* 1 × 10^*−*12^), resulting in significant localization *p*-values (*p* = 9.89 × 10^*−*3^ and 2.48 × 10^*−*2^, respectively). In contrast, *H19* and *Nnat* show weak or non-significant depletion (depletion *p* = 0.140 and 0.103), and consequently do not achieve significant localization (localization *p* = 0.089 and 0.082; Fig. 1).

We provide a secondary set of examples using a widely used 10X PBMC dataset [8] to reinforce these concepts in Supplementary Fig. S2.

### 2.2 Locat shows high sensitivity and robustness in synthetic data experiments

To characterize how Locat behaves under controlled conditions, we generated synthetic datasets with known ground truth and evaluated its response to changes in sample size, sparsity, and localization radius. Each simulation consisted of a unit-variance multivariate Gaussian background distribution of 1500 cells, with expression for a simulated localized gene drawn from a compact neighborhood (red points, Supplementary Fig. S3A, C, and E). We show three statistical approaches for testing simulated genes: Locat’s localization *p*-value (see Section 4.1.5), a likelihood ratio test comparing gene-weighted versus background density models (see Section 4.6), and a permutation-based variant of Locat (see Section 4.7). The likelihood ratio test serves as a natural parametric baseline that directly compares distributional fits but lacks explicit constraints for compactness or specificity. The permutation-based approach is a resampling method that estimates significance directly from randomized replicates of the data rather than from a parametric null; it is often favored because it makes minimal distributional assumptions, but it incurs prohibitive computational cost and, as the results below show, is not well calibrated in this setting. For each experimental setup, 50 replicate trials were performed.

#### Sensitivity to sample size

We varied the number of expressing cells between 25 and 251 while maintaining localization around a fixed center (Supplementary Fig. S3A). Locat’s localization *p*-value remained consistently below the significance threshold *α* = 0.05 for much of the data, indicating high sensitivity even for genes with small sample sizes (Supplementary Fig. S3B, left panel). In contrast, the likelihood ratio test performed poorly, with sensitivity decreasing as the number of expressing cells increased (Supplementary Fig. S3B, middle panel). This reflects increasing overlap between the gene and background densities as expression becomes more widespread. The *p*-values from the permutation-based approach indicated that the approach yielded similar sensitivity to Locat’s localization test but exhibited greater variability in *p*-values at larger sample sizes due to the finite number of resampling iterations (Supplementary Fig. S3B, right panel). These results show that Locat maintains high sensitivity across a large range of gene-expressing population sizes, spanning simulated genes expressed in 1.6% to 16% of the total number of cells in our simulation.

#### Robustness to gene sparsity

In our second simulation scenario we introduced a multimodal localized population with increasing positional jitter with respect to each mode, to represent different levels of compactness in a local support region. Locat is intended to identify extremely compact genes as well as genes that are moderately compact but very specific to a particular population of cells. This experiment evaluated simulated genes with progressively more spread out expression patterns, representing increasing gene sparsity in the expressed population. The simulated genes are also multimodal, which is an additional source of deviance from the background distribution (Supplementary Fig. S3C). The likelihood ratio test lost power rapidly, with *p*-values exceeding the significance threshold already at a jitter of 5% (Supplementary Fig. S3D, middle panel). In contrast, Locat’s localization *p*-value remained significant across most of the tested jitter range and only lost significance once the jitter was sufficiently large that the expression pattern became indistinguishable from the background distribution (Supplementary Fig. S3D, left panel). The permutation-based test maintained nominal significance across all jitter levels, including at extreme dispersions, suggesting overly permissive behavior when gene expression structure is nearly indistinguishable from the background signature. (Supplementary Fig. S3D, right panel).

#### Sensitivity to localization radius

In our third simulation scenario we varied the dispersion radius of the localized expression region while holding the background variance constant (Supplementary Fig. S3E). As the dispersion radius increases, the signal gradually merges with the background distribution, thus losing its localization property. Modeling concentration as a naïve single-sided z-test we expected no identifiable localization for a dispersion radius greater than 1.6 at a significance threshold of *α* = 0.05, i.e. the observed distribution of the gene expression becomes undistinguishable from the distribution of the background. Locat’s localization *p*-value crossed the significance threshold *α* = 0.05 at dispersion radius around 1.6. The likelihood-ratio test lost significance by radius ≈ 0.5 (Supplementary Fig. S3F, left and middle panels). The permutation-based approach remained significant across nearly all radii, confirming its anti-conservative tendency and indicating a greater tendency toward false positives compared to Locat (Supplementary Fig. S3F, right panel). These findings indicate that Locat yields higher sensitivity than the likelihood-ratio test approach and higher specificity than the permutation-based approach.

These simulations demonstrate that Locat maintains power across rare populations (≥ 1.6% prevalence), handles multimodal distributions and positional noise, and correctly fails to detect localization when the signal concentration equals the background. The latter property is critical for non-uniform embeddings e.g., single-cell RNA data, where likelihood ratio and permutation tests show systematic biases.

### 2.3 Comparison of marker gene selection approaches

We compared the performance of Locat against existing methods on the task of identifying localized genes. Evaluating this task is complicated by the absence of an established gold standard or benchmark dataset for gene expression localization, i.e., expression that is enriched within a compact subset of cells, together with marked depletion elsewhere. Existing marker gene resources such as CellMarker provide curated associations between genes and particular cell types, but do not require a marker to exhibit depletion outside its associated population. Such resources therefore do not supply ground truth for the stricter, depletion-aware notion of localization considered here.

Rather than relying on any curated list of marker genes, we instead used the cell-type partition itself, independent of its associated gene annotations, as a reference structure against which to measure specificity. Cell typing or gene expression clustering partitions the expression manifold into groups of cells with similar overall expression profiles; this grouping, rather than any predefined marker set, is what we use to ask whether a given gene’s expression is concentrated within one such group and depleted across the rest, which is the property our definition of localization requires. The population associated with a localized gene need not correspond exactly to any single predefined group, since localized expression may occur within a subset of a group or span cells belonging to several groups; nevertheless, because these groupings capture the major distinctions among populations of similar cells, we expect genes with genuinely localized expression to exhibit high specificity to a single group, while genes whose expression is dispersed across several distinct populations will exhibit correspondingly lower specificity. This group specificity therefore offers a fair and interpretable basis for comparing methods on the localization criterion we adopt, without relying on a predefined, and necessarily incomplete, set of ground-truth localized markers.

To quantify this, we used the cell-type specificity score *S*_*g*_ (Methods 4.8, Eq. 30), applied to annotated cell types or dominant clusters as appropriate. *S*_*g*_ is the maximum proportion of a gene’s total mean expression attributable to any single group, with higher values indicating greater specificity. We compared the mean *S*_*g*_ among the top-ranked genes returned by each method to assess the extent to which each approach prioritizes genes with expression specific to a single cell type or dominant cluster over genes with expression broadly distributed across these groups.

We evaluated cell type specificity in top genes returned by several approaches, enabling us to compare Locat to diverse frameworks: LMD (diffusion statistic over a cell-cell graph, [5]), singleCellHaystack (2D kernel density comparison, [2]), Hotspot (graph autocorrelation based on Geary’s C, [3]), GSPA (gene-specific pattern analysis, [6]), GiniClust (Gini index-based rare cell detection, [4]), and Scanpy rank genes groups (Wilcoxon rank-sum test on user-defined groups, [9]). We also included a spectral baseline (SpectralRH) that scores genes by graph smoothness and concentration of spectral energy in low-frequency Laplacian eigenmodes on a k-nearest-neighbor graph and serves as a comparator with a different objective from the other methods (see Supplementary Methods S1.8.2). Several of these methods are designed explicitly to capture local clustering patterns, whether through autocorrelation (Hotspot) or diffusion-based statistics (LMD). Whereas the other approaches prioritize compactness, local autocorrelation, or slow diffusion across neighboring cells, SpectralRH is intended to highlight genes whose expression aligns with well-supported transitions and boundaries in the embedding (see Supplementary Methods S1.8.1, S1.8.2).

Our cell type specificity analyses examined the mean cell type specificity observed in the top 100 genes recovered by each of the marker selection method. We performed this same comparison in four primary datasets, including murine embryonic dermis, two examples of normal peripheral blood mononuclear cells (PBMCs), and IFN-*β*-stimulated PBMC datasets. In these data, Locat consistently recovered genes with higher cell type specificity than genes recovered by most or all other methods (Fig. 2A–D). This advantage also extended to three additional datasets, including *Chlamydomonas reinhardtii* transcriptomes under iron-sufficient and iron-deficient conditions [10] and murine bone marrow from the Tabula Muris Senis consortium [11] (Supplementary Fig. S8). Qualitative inspection of top-10 gene expression patterns on the UMAP embedding confirmed that Locat consistently selected genes with visibly constrained background expression across these datasets (Supplementary Figs. S9– S11). This finding suggests Locat is able to prioritize genes that are localized to particular cell populations and largely absent/depleted from other populations. Next to Locat, the most competitive methods were GSPA and SpectralRH, which showed similar performance in certain datasets (Fig. 2A, 2C). In The 3k PBMC cell dataset, the evaluated methods showed lower overall specificity of top genes for all methods (Fig. 2C) as compared to the other datasets evaluated. In this dataset, GSPA had slightly greater cell type specificity than Locat with substantial overlap in confidence intervals. Lower cell type specificity in this data suggests that localized genes in this dataset tended to bridge multiple cell types, making it more difficult to distinguish between population-specific genes and diffuse genes by cell type specificity scores alone. To further examine differences in top genes selected by different approaches, we compared rankings of individual genes in the murine embryonic dermis dataset [7]. This dataset contains approximately 5000 cells spanning the transition from upper dermal fibroblasts to *Sox2* -positive dermal condensates, early precursors of hair follicles. This setting presents a challenging test case: it contains two intersecting sources of variation. There is continual progression of fibroblasts toward dermal condensate identity, and these fibroblasts likewise move through phases of the cell cycle. Together, these form a continuous transcriptional manifold with continuous cluster boundaries (Supplementary Fig. S4A). This comparison uses the same set of methods introduced above. Although many of the marker genes prioritized by Locat and other approaches were relatively cell-type specific, the continuous or fuzzy cluster boundaries of this dataset presents a challenging case for traditional cluster-based differential expression methods, motivating the need for clustering-free marker gene selection approaches.

**Fig. 2.**
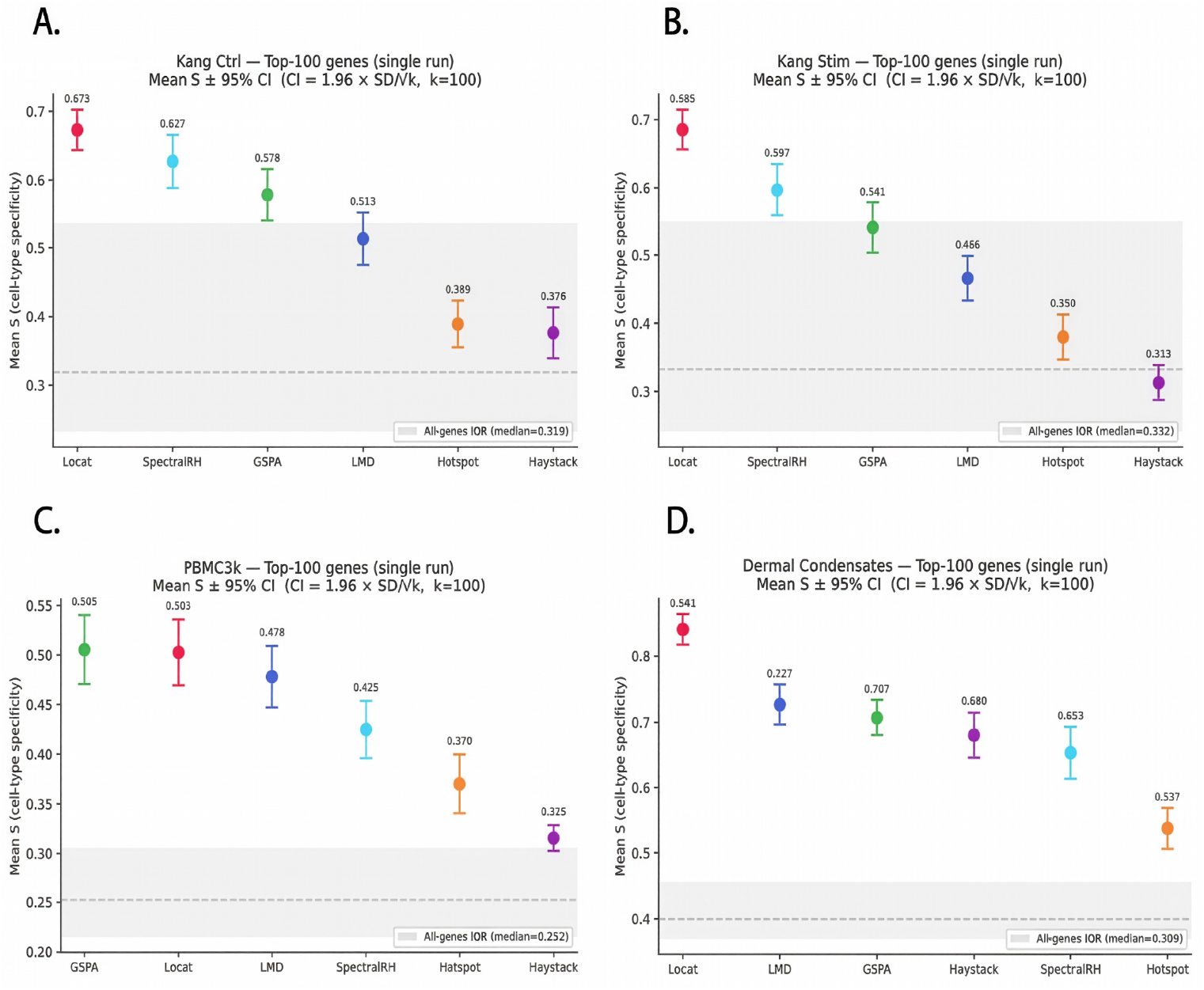
Locat achieves higher cell-type specificity than competing methods in several datasets. Mean cell-type specificity score *S* (maximum proportion of total expression attributable to any single cell type) *±* 95% CI 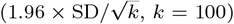for the top-100 genes returned by each method. Methods are ordered left to right by decreasing mean *S* within each panel. Gray shaded region: interquartile range of *S* across all genes; dashed line: all-genes median *S*. **A**. Kang control PBMCs. **B**. Kang IFN-*β*-stimulated PBMCs. **C**. 3k PBMC dataset. **D**. Murine embryonic dermis. Locat ranks first in mean cell-type specificity in three of four datasets; in the 3k PBMC dataset (C), Locat and GSPA achieve comparable scores with overlapping confidence intervals, and all top methods show lower absolute specificity, consistent with a dataset in which marker genes frequently bridge multiple cell populations.

Genes prioritized by Locat, including *Hist1h2bb* (Locat: 98th percentile; LMD: 94th percentile, singleCellHaystack 73rd percentile), *Casq1* (Locat: 99th percentile; GSPA: 87th percentile; LMD: 95th percentile, Hotspot: 64th percentile), and *Qprt* (Locat: 99th percentile; GSPA: 91st percentile; LMD: 96th percentile; Hotspot: 74^th^ percentile), exhibited tightly confined expression with minimal background (Supplementary Fig. S5). In contrast, genes ranked more highly by enrichment-focused methods, including *Col8a1* (LMD: 93rd percentile; GSPA: 99th percentile, Locat: 76^th^ percentile) and *Ptn* (LMD: 99th percentile, SpectralRH: 98th percentile; singleCell-Haystack: 99th percentile; Locat: 5th percentile) showed moderate local enrichment accompanied by substantial background expression (Supplementary Fig. S5). Such diffuse patterns achieve high scores under concentration-only criteria and lack well-defined localized domains, otherwise enforced by the depletion score included in Locat. We provide additional visualizations of genes with high rank discrepancies between Locat and existing models in Supplementary Fig. S6.

Rank correlation analysis across all genes revealed low to moderate concordance between Locat and alternative approaches (Fig. 3A). Locat shows highest concordance with GSPA and LMD (Spearman *ρ* = 0.66 and *ρ* = 0.63, respectively) and moderate agreement with SpectralRH (*ρ* = 0.38), reflecting a shared emphasis on detecting genes with highly concentrated expression, though Locat’s dual concentration-depletion framework further penalized genes with broad and nonspecific background expression, as evidenced by Supplementary Fig. S6. We additionally computed Jaccard indices between the top 500 genes identified by each method, and evaluated rank correlations between methods restricted to the top 500 genes identified by Locat (Supplementary Fig. S7). In both analyses, concordance remained modest (Supplementary Fig. S7). This indicates that the depletion test introduces a distinct selectivity criterion that separates truly localized markers from broadly expressed genes with local peaks. The rank- and overlap-based comparisons in this section establish that Locat prioritizes a distinct class of genes relative to existing approaches. The following sections provide broader context for these differences through applications in dermal condensate differentiation (Section 2.5), interferon-*β* stimulation (Section 2.7), ESC differentiation time series (Section 2.8), and controlled perturbations of a real-data template gene (Section 2.4).

**Fig. 3.**
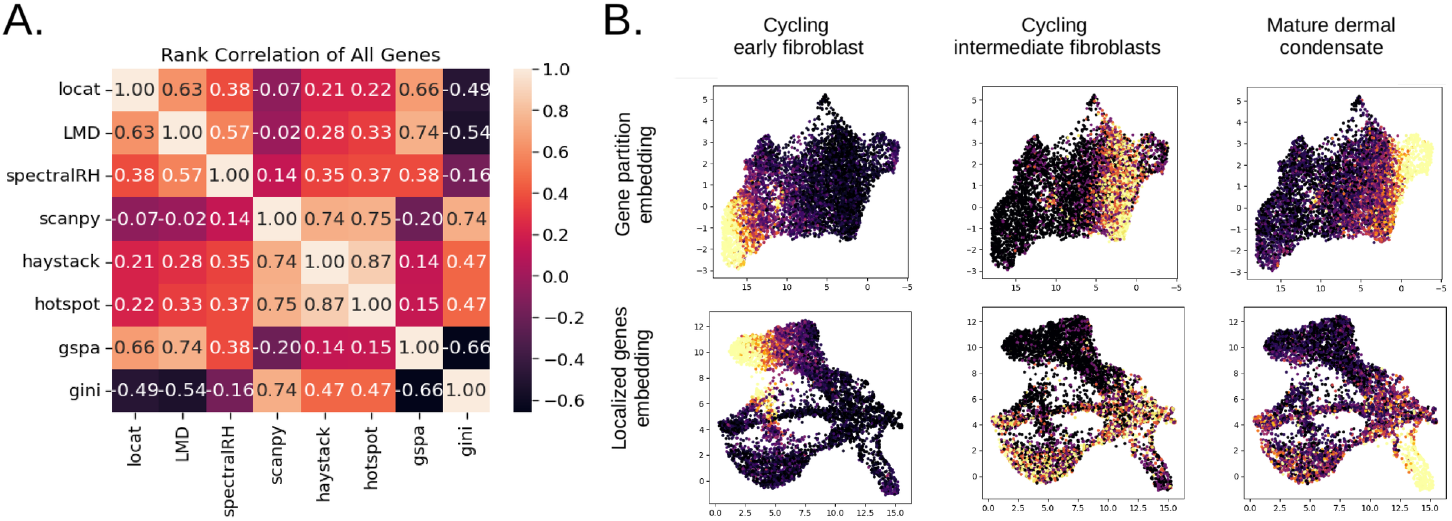
Locat identifies distinct gene programs and shows moderate concordance with alternative prioritization methods. **A**. Spearman rank correlation heatmap across eight gene prioritization methods applied to the murine embryonic dermis dataset. **B**. Average expression of three Locat-derived modules (cycling early fibroblasts, cycling intermediate fibroblasts, mature post-mitotic dermal condensates) identified from a differentiation-focused gene partition [16]. Top row: module expression on the differentiation-focused UMAP, illustrating alignment with the fibroblast-to-dermal-condensate trajectory. Bottom row: the same modules on the localized-gene embedding, where each module occupies a spatially restricted region. Additional module views are in Supplementary Fig. S17.

### 2.4 Comparing methods using simulated perturbations of a real dataset

To determine whether Locat’s distinct gene selections reflect a meaningful advantage rather than merely a different ranking criterion, we performed controlled gene-level simulations to evaluate how different gene prioritization methods respond to simulated genes with variable sample size and background expression. Simulations were based on the murine dermis dataset, using *Sox2* as a template gene due to its compact and well-defined expression pattern in dermal condensates. Synthetic variants were generated by perturbing *Sox2* expression while preserving the underlying cell embedding.

Each simulation retained the original cell embedding and expression values for approximately 1000 randomly sampled background genes, together with multiple simulated copies of *Sox2*. This experiment was conducted independently of the full-dataset correlation analysis and isolates method behavior under controlled perturbations of a single gene. For methods that do not report *p*-values, including GSPA and LMD, significance thresholds are defined empirically from the distribution of gene scores (see Section 4.8.1). All methods introduced above were applied identically in this setting.

#### Robustness to subsampling

We assessed robustness to decreasing prevalence by progressively subsampling the number of cells expressing the simulated *Sox2* gene while maintaining its expression pattern (Supplementary Fig. S12A). Locat maintained detectability across nearly the full range of subsampling levels (Supplementary Fig. S12B). SpectralRH, Hotspot, and singleCellHaystack similarly showed limited sensitivity to subsampling in this regime. By contrast, cluster-based differential expression tests showed a rapid sensitivity drop as expressing cells became rare. GSPA and LMD were significant at some moderate subsampling levels, but once more than half of expressing cells were removed, both showed a monotonic loss of significance with further subsampling.

#### Sensitivity to background expression

We next evaluated sensitivity to loss of localization by progressively redistributing *Sox2* expression into the background population while holding the number of expressing cells fixed (Supplementary Fig. S13A). In this setting, Locat *p*-values became less significant and crossed the detection threshold as expression became broadly distributed and indistinguishable from background (Supplementary Fig. S13B). SpectralRH, GSPA, and LMD similarly exhibited monotonic score increases with increasing background expression, indicating these methods were also sensitive to this attribute. In contrast, Hotspot, singleCellHaystack, and cluster-based differential testing remained largely insensitive until background expression signal became extreme, at which point they no longer detected the simulated genes.

These simulations, based on controlled perturbations of a single template gene (*Sox2*) in one dataset (murine embryonic dermis), indicate that among the tested methods, Locat alone balances two competing but essential criteria for marker discovery: robustness to decreasing gene prevalence and sensitivity to loss of localization due to background expression. While methods such as LMD, GSPA, and SpectralRH also penalize increasing background expression, those same methods exhibited unstable or rapidly degrading behavior under subsampling, indicating that their apparent sensitivity to background signal is not preserved when expression becomes sparse. Conversely, methods that remained robust to subsampling largely failed to penalize background expression, continuing to assign high confidence to genes whose expression was broadly distributed rather than localized. We treat this as evidence from a single experiment, not a claim established across all genes or biological systems more broadly.

This trade-off has direct implications for real data analysis. Some genes are expressed in many cells or show strong local peaks, yet still remain broadly distributed across the embedding. Methods that do not adequately distinguish this pattern may rank such genes highly even when they are not specific markers of a localized cell state. Locat is better able to separate these genes from genes whose expression is concentrated within a restricted region of the embedding, even when those localized genes are present in relatively few cells. As a result, gene prioritization depends more strongly on spatial specificity than on prevalence or expression magnitude alone.

### 2.5 Localized markers recover major axes of biological variation

#### Localized markers along developmental and cell-cycle axes

In the murine embryonic dermis dataset (see 2.3) Locat identified 430 out of 11700 genes at *p <* 0.05 (calibration of this threshold under simulation-based null analyses is shown in Supplementary Fig. S1; see Supplementary Methods S1.3) with compact expression patterns aligned with both cell-cycle progression and differentiation-related variation. Several cell-cycle-associated genes including *Hist1h2bb* (*p* = 2.34 ×10^*−*2^),*Ccnb1* (*p* = 4.27 ×10^*−*2^), and *Kif20a* (*p* = 4.90 ×10^*−*2^) formed sharply confined domains corresponding to distinct cell-cycle phases, consistent with established phase-specific programs [12]. Genes associated with dermal condensate identity, including *Sox2* (*p* = 1.05 ×10^*−*2^), *Foxd1* (*p* = 1.97 ×10^*−*2^), and *Gal* (*p* = 1.96 ×10^*−*2^) localized to terminal regions of the fibroblast-to-dermal-condensate trajectory, in agreement with their known expression patterns in developing skin [13, 14]. Several genes linked to ephrin signaling and cytoskeletal dynamics localized to a cycling population adjacent to the terminal dermal condensate (Supplementary Fig. S14). These included *Epha8* (*p* = 3.49 ×10^*−*3^) and *Stmn2* (*p* = 4.48 ×10^*−*2^). This pattern is consistent with a transitional population undergoing cytoskeletal and signaling remodeling prior to terminal condensate differentiation [15].

#### Manifold reconstruction from localized genes

To test whether Locat-prioritized genes retain biologically meaningful structure in the dataset, we constructed a localized-gene embedding (LG-embedding): a low-dimensional representation computed using only the 430 significant localized genes discovered by Locat (Supplementary Fig. S15). We compared this to a baseline UMAP embedding derived from 4000 highly variable genes (HVG-embedding, Supplementary Fig. S4). Despite using substantially fewer genes, the LG-embedding recapitulated both the fibroblast-to-dermal-condensate differentiation trajectory and the cell-cycle–associated variation observed in the HVG embedding (Supplementary Fig. S15). These results indicate that genes selected for compact localization are sufficient to represent at least two well-characterized axes of biological variation in this system, demonstrating that localized markers, though individually restricted to confined expression domains, can collectively preserve continuous developmental and cell-cycle structure using a substantially smaller and more interpretable feature set than the HVG baseline.

### 2.6 Localized markers reveal distinct stages of development in murine embryonic dermis

#### Module discovery reveals cell-cycle exit and lineage commitment

A key question in dermal development is identifying the molecular signatures that mark fibroblast commitment to the dermal condensate lineage and the timing of cell-cycle exit during this transition. To address this, we constructed a differentiation-focused embedding from the murine embryonic dermis dataset (see 2.3) by removing cell-cycle variation using gene partitions from [16] (Supplementary Fig. S16). We then applied Locat to identify which genes from the entire dataset were localized on this new embedding. By construction, such localized genes along the developmental trajectory correspond to specific developmental stages.

Locat identified 1302 distinct localized genes along the developmental trajectory, which we grouped in 8 gene modules (see 4.2). These modules exhibited particular expression patterns corresponding to early fibroblast, intermediate, and mature dermal condensate states (Fig. 3B, upper row). When we projected these module expression signatures back onto the LG-embedding (which retains cell-cycle structure, Fig. 3B, lower row), the modules traced a continuous developmental path revealing discrete transitions from proliferative to post-mitotic states, highlighting a region of the embedding associated with cell-cycle exit and dermal condensate identity (Fig. 3B, right column).

This analysis highlights a key feature of Locat: because it is embedding-agnostic (see Section 4.1.1), it can be applied to any user-defined low-dimensional representation and will identify genes localized with respect to that representation. The LG-embedding derived from all localized genes captured both developmental and cell-cycle variation (Supplementary Fig. S15). When an embedding is constructed to emphasize a single biological axis, such as differentiation, applying Locat to that representation prioritizes gene programs whose expression patterns align with the chosen trajectory. In this setting, the resulting modules reflect genes whose localization is most consistent with the differentiation-focused embedding, reducing contributions from other sources of variation (e.g., cell-cycle state) to the extent that they are not encoded in that representation.

### 2.7 Single-sample localization analysis avoids integration artifacts in multi-condition experiments

Multi-sample single-cell studies commonly integrate datasets using batch-correction methods that align cells from different conditions into a shared representation. These approaches reduce cross-sample variation to facilitate direct comparison of similar cell states. However, not all cross-sample differences reflect technical noise. In many settings, they represent true biological programs induced by perturbation, genotype, or developmental state. Because batch correction explicitly minimizes between-sample variation, it can attenuate condition-specific expression structure that is often the primary focus of analysis.

We therefore take a different approach. Rather than applying batch-correction–based integration to align cells across conditions into a shared embedding prior to analysis, we apply Locat independently within each sample and evaluate gene localization with respect to that sample’s native representation. Localization scores can then be compared directly across samples, or cells from multiple samples can be embedded together using the union of condition-specific localized genes as the feature set. In this setting, a combined embedding is constructed without additional batch correction, using only genes identified as localized. This strategy preserves condition-specific structure during localization analysis while still enabling interpretable cross-condition comparison at the level of gene programs. Such combined embeddings are most appropriate when samples share a common biological architecture but differ due to defined perturbations (e.g., treatment versus control or genotype), so that a shared representation remains meaningful. When samples differ more fundamentally (e.g., across developmental time points, tissues, or technologies), cross-sample embedding may be less informative. In these settings, localization patterns are more appropriately interpreted within each sample.

#### Integration artifacts obscure stimulus-specific localized programs

We demonstrate this framework using a single-cell RNA interferon-*β* stimulation dataset [17], comprising approximately 12000 peripheral blood mononuclear cells per condition from a single donor. Control and IFN-*β*-stimulated samples were processed independently: each was normalized separately, embedded in its own PCA space, and analyzed with Locat to identify localized genes without cross-sample alignment. The two samples were subsequently merged into a combined dataset of approximately 24000 cells for visualization and comparison. This design avoids imposing batch-correction-based alignment during localization analysis.

Given that the two samples originate from the same donor, an embedding constructed from both control and stimulated cells should align shared immune populations while preserving stimulus-induced changes within responsive cell types. We therefore evaluated whether merging the datasets could achieve both goals, either without batch correction or after applying batch correction. A UMAP embedding computed from highly variable genes on the merged data without batch correction showed near-complete separation of control and stimulated cells (Fig. 4A). This separation likely reflects technical differences between samples more than the biological effects of IFN stimulation and prevents meaningful comparison of shared immune populations. Applying Harmony-based batch correction [18] substantially improved cross-condition alignment (Fig. 4B), but visual inspection and differential abundance analysis (Supplementary Fig. S18) indicate that stimulated monocytes no longer form a distinct region in the embedding. Monocytes are expected to mount a robust interferon response characterized by induction of canonical interferon-stimulated genes and inflammatory mediators, including chemokines and antigen-presentation programs [19], which should generate a transcriptionally distinct stimulated state relative to control. In the Harmony-corrected embedding, this separation is no longer apparent. These results highlight a central challenge: merging samples without batch correction exaggerates technical differences, whereas strong batch correction can obscure biologically meaningful stimulus-specific structure.

**Fig. 4.**
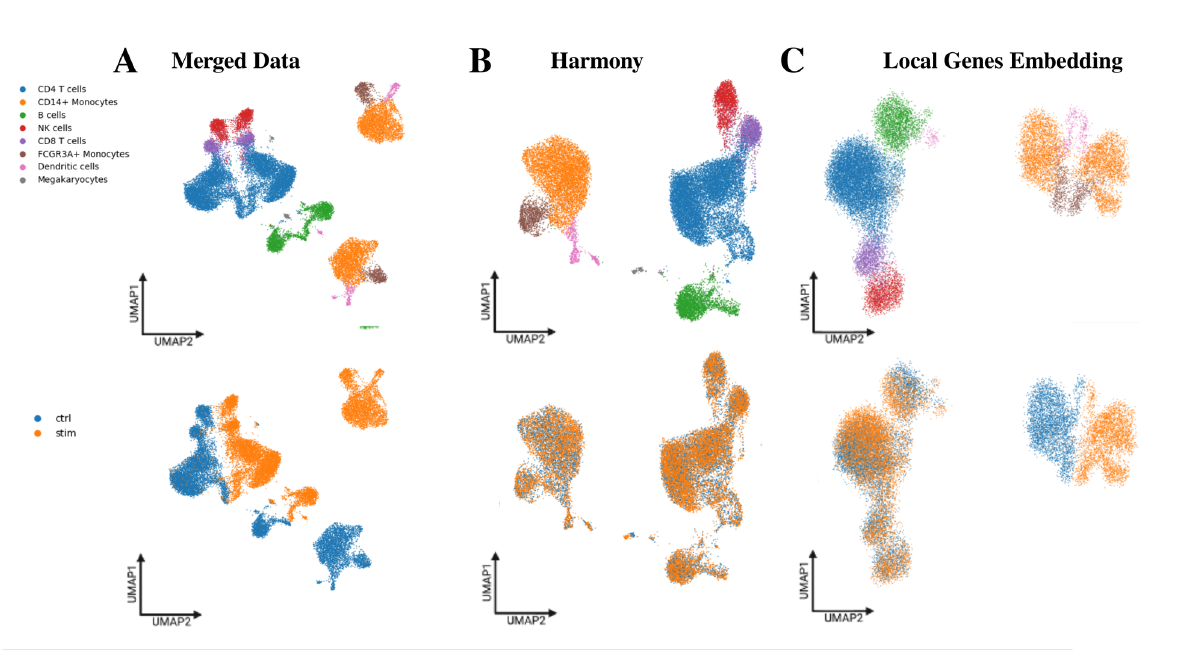
Locat identifies condition-specific localized gene programs in an informative embedding of IFN-*β*–stimulated and control PBMCs (A–C) Locat distinguishes shared and stimulus-specific transcriptional programs without batch correction. (A) UMAP of merged control and IFN-*β* samples showing complete separation before correction. (B) Harmony-corrected embedding with over-alignment and loss of biological signal. (C) Embedding based on top localized Locat genes preserves shared cell types while maintaining stimulus-specific separation of monocytes.

As an alternative approach to representing both samples jointly, we generated a UMAP embedding using only genes that were significantly localized in at least one condition (1283 genes from control, 1398 genes from stimulated, 944 genes shared). Under our dual concentration–depletion criterion, these genes serve as discriminative features because their expression is confined to well-defined cell populations rather than broadly distributed across the manifold. The resulting embedding aligned shared immune lineages while preserving a distinct region corresponding to IFN-*β*-responsive monocytes, without requiring batch correction (Fig. 4C).

We then used this co-embedding to visualize genes exhibiting differential localization between conditions. For validation, we also plotted expression on embeddings constructed separately for the control and stimulated samples to confirm that specific genes were localized uniquely within IFN-*β*-responsive monocytes or within control monocyte populations.

#### Condition-specific differential localization

We define differential localization as a gene that is significantly localized in one condition but not significantly localized in the other. We identified such genes by comparing Locat’s localization *p*-values across the two samples using a fixed significance threshold. Using this criterion, 454 genes were significantly localized in the stimulated sample but not in control, and 339 genes were significantly localized in control but not in stimulated.

Genes with much stronger localization in the stimulated sample included *CCNA1, CXCL9, NUPR1, CH25H*, and *JUP*, which were confined to interferon-responsive monocytes (Supplementary Fig. S19A and Supplementary Fig. S20A). Among these, *CXCL9* and *CH25H* are well-established interferon-stimulated genes involved in chemokine signaling and antiviral lipid metabolism, respectively [20, 21]. *NUPR1* reflects stress- and interferon-associated transcriptional responses in activated myeloid cells [17, 22].

In contrast, several genes were significantly localized in the control condition but not in the stimulated condition, including *TPST1, MMP9, NAB2, OSM*, and *HIPK2* (Supplementary Fig. S19B and Supplementary Fig. S20B). *MMP9*, which has been associated with extracellular matrix remodeling and basal monocyte function [23], marked a transcriptionally restricted subset of unstimulated monocytes. *OSM*, a cytokine with context-dependent roles in myeloid cells [24], was likewise confined to a subset of control monocytes. Regulatory genes such as *NAB2*, a transcriptional regulator [25], and *HIPK2*, a kinase involved in transcriptional control and stress signaling [26], were similarly localized specifically in control samples. The loss of localized expression for these genes following interferon stimulation suggests that these restricted baseline programs are disrupted under type I interferon exposure, consistent with the well-established reprogramming of monocyte transcriptional states in response to interferon signaling [19, 27].

These results indicate that single-sample localization analysis, followed by joint visualization using condition-specific localized genes, preserves biologically interpretable stimulus-associated programs that may be obscured by integration-based approaches.

#### Application to subsequent analyses

This principle of independent per–sample Locat analysis followed by comparison of localization patterns across samples, or localized-gene-based joint visualization when needed, is applied throughout subsequent analyses (see Section 2.8). By avoiding premature integration, we preserve condition-specific localized structure in expression space that would otherwise be erased by alignment procedures. This strategy is particularly critical for detecting rare cell states, transient activation programs, and responses specific to the one condition or sample group in multi-sample experimental designs.

### 2.8 Temporal localization dynamics reveal stage-specific regulatory programs during ESC differentiation

Having established the principle of independent per-sample localization analysis (see Section 2.7), we applied this framework to characterize temporal dynamics of gene localization during cellular differentiation. We analyzed a retinoic acid (RA) differentiation time course profiling mouse embryonic stem cells (ESC) at days 0, 2, 4, and 10 following induction, with a total of 8 samples, each representing a biological replicate of one time point [28]. This dataset has been previously used to define lineage-specific regulatory programs across developmental time [28]. We asked: (i) How do localized gene sets change across developmental stages? (ii) Are localization patterns reproducible across independently analyzed replicates? (iii) Can temporal trajectories of gene localization identify stage-specific regulatory programs? (iv) Can localized genes be used to anchor specific cell populations between samples and time-points?

Each sample was analyzed independently with Locat using the sample’s own PCA embedding without batch correction, thus preserving sample-specific embedding structure while enabling comparison of localization scores across developmental stages (Fig. 5). Unless otherwise specified, localization at a given timepoint was determined by comparison of the localization *p*-values of the given timepoint, combined using geometric mean, to the significance threshold.

**Fig. 5.**
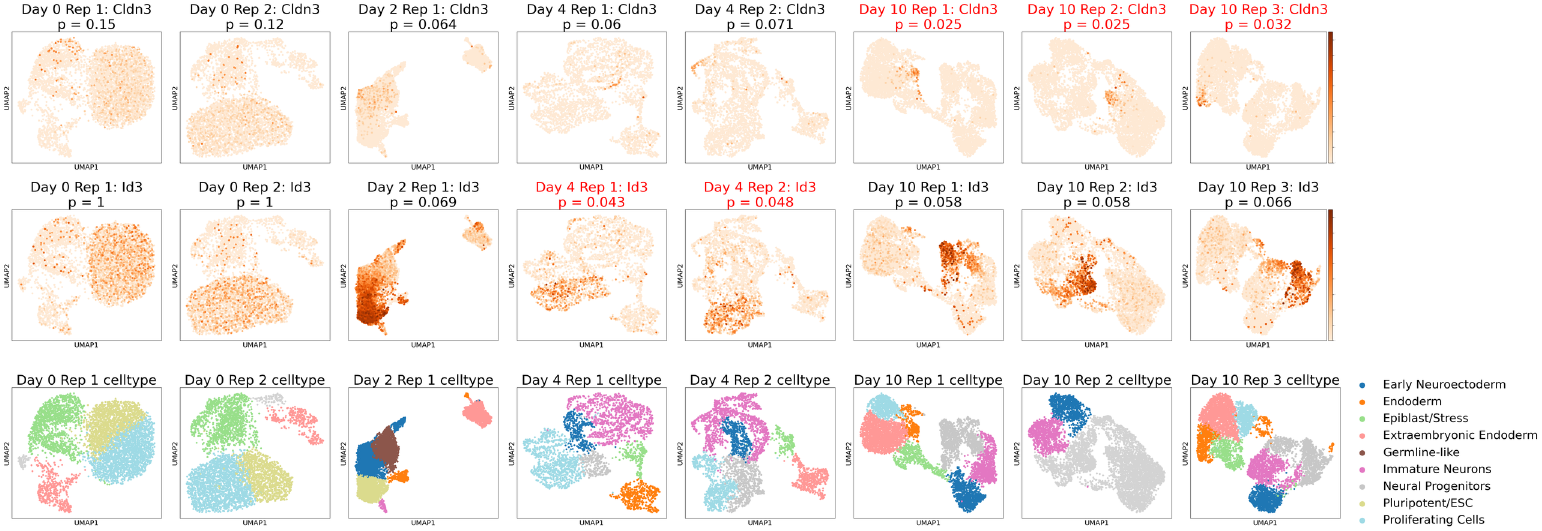
Locat reveals dynamic localization patterns in RA-induced murine ESC developmental timecourse. UMAP embeddings for eight samples from day 0 to day 10 of RA treatment. Top and middle rows plot average expression of two genes that are conditionally localized in phases of differentiation after RA exposure. Titles in the first two rows track magnitude and significance of localization *p*-values. The third row displays annotated cell types corresponding to each of the eight samples.

#### Progressive increase in localization marks neural differentiation

Locat identified 2431 genes as significantly localized (*p <* 0.05) at at least one time-point. The number of localized genes increased across the time course: 432 genes at day 0, 515 at day 2, 1323 at day 4, and 1429 at day 10. Full gene-level results are provided in the Supplementary Data CSV. This increase reflects progressive concentration of gene expression into restricted neighborhoods in latent expression space as differentiation proceeds, consistent with the emergence of increasingly distinct cellular states from initially more homogeneous progenitor populations.

To identify coordinated temporal patterns, we assigned each gene from the localized gene set to one of the 15 possible temporal localization clusters representing the time points in which the gene showed significant localization (by construction we excluded the case in which the gene was not localized in any timepoint). Chi-square analysis revealed highly significant non-independence of temporal localization patterns (*p* ∼2.51 ×10^*−*150^), indicating that localization did not occur randomly but exhibit structured temporal dynamics. The observed temporal structure is likely reflecting the dynamics of biological processes, as supported by our analyses below.

#### Five dominant temporal patterns capture developmental regulation

We grouped genes according to their temporal localization patterns across time points and quantified over-representation for each pattern using *δ*, defined as the difference between the observed frequency of genes assigned to that pattern and the frequency expected if temporal pattern membership were independent across time points. To assess whether observed deviations exceeded what would arise by chance under this independence assumption, we estimated the null distribution of *δ* using Monte Carlo simulations (*n* = 10000). Under this null model, the mean deviation was 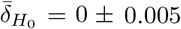, and the largest deviation observed was max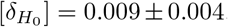. We identified five temporal patterns whose observed *δ* values substantially exceeded this null range (Supplementary Fig. S21; cluster-level *δ* values are summarized in Supplementary Table S2 and the full gene-level results are provided in the Supplementary Data CSV).

1. **Specific emergence on day 10** (cluster 0001, *n* = 818, *δ* = 0.090). The most over-represented pattern contained genes whose localization became detectable only at day 10, reflecting late-emerging, highly specific expression during later stages of differentiation. Localized genes in this group included regulators associated with ventral neural patterning and lineage specification, including *Nkx2-2* and *Gli2* [29–31], as well as membrane and signaling genes associated with differentiated glial states such as *Gpr17* [32] and *Cldn11* [33]. These 818 genes showed a progressive increase in mean localization scores across the time-course (negative log-10 *p*-values: 0.93 on day 0, 1.07 on day 2, 1.14 on day 4, 1.42 on day 10), indicating moderate concentration and moderate depletion that strengthened into significant localization only at terminal differentiation. This pattern suggests that lineage-specific programs begin subtly and strengthen through clonal expansion, migration, or selective survival rather than arising de novo at late stages. As a result, while differentiation progresses, cell types become more segregated on the embedding due to increasingly stronger diversification of transcriptional programs.
2. **Specific transient localization on day 4** (cluster 0010, *n* = 624,*δ* = 0.049). This cluster captured genes localized at day 4 following retinoic acid exposure, reflecting a short-lived reorganization of transcriptional programs during early post-induction responses. Localized genes in this cluster included factors associated with pluripotency exit and early differentiation, including *Tfcp2l1, Zfp42, Tbx3*, and *Klf5* [34], together with epithelial-associated genes such as *Krt8, Epcam*, and *Esrp2* [35, 36]. The coexistence of pluripotency-linked and epithelial organizational genes suggests that this pattern reflects a transient intermediate state during the early response to retinoic acid, before stabilization of more restricted lineage-specific transcriptional programs [34, 37].
3. **Specific transient localization on day 2** (cluster 0100, *n* = 223, *δ* = 0.039). Genes in this cluster localized specifically at day 2, defining an early and transient pattern that coincides with pluripotency exit and the beginning of lineage specification. Localized genes in this cluster included regulators associated with pluripotency and its early dissolution, such as *Nanog, Zscan4c*, and *Esrrb* [38–40], as well as early neural and patterning factors including *Sox21* and *Pax3* [29]. This cluster also included epithelial-associated genes such as *Cldn2* and *Aqp3* [35]. The restriction of localization to day 2 indicates that these programs are transiently organized during an intermediate differentiation stage, after pluripotency exit but before stable lineage commitment [34].
4. **Specific localization at day 0** (cluster 1000, *n* = 199, *δ* = 0.038). Genes in this cluster were localized at day 0 prior to induction, reflecting a baseline cell state organization characteristic of the pre-differentiation condition that is rapidly altered upon retinoic acid exposure. Localized genes in this cluster included transcriptional regulators associated with pluripotency maintenance and early developmental programs, such as *Prdm14* [41], *Zbtb16* [42], and *Meis3* [43], alongside genes involved in cell–cell adhesion, extracellular matrix organization, and baseline cellular structure, including *Cdh6, Cdh9, Dcn*, and *Bgn*, consistent with stable regulatory and structural programs in pluripotent or pre-induction cell states [43, 44]. This cluster also contained genes linked to homeostatic signaling and surface or receptor-associated functions, such as *Thy1, Il15ra*, and *Trem2*, consistent with transcriptionally stable but heterogeneous baseline populations [45]. The restriction of localization to day 0 suggests that these programs are specific to the pre-induction state and are rapidly dissolved or reorganized upon retinoic acid exposure, reflecting loss of the initial baseline configuration rather than persistence of a differentiated identity [46].
5. **Sustained localization: days 0** →**2** →**4** (cluster 1110, *n* = 47, *δ* = 0.009).Genes in this cluster exhibited sustained localization from day 0 through day 4, but not at day 10, indicating persistent cell populations during early differentiation prior to later state diversification. This set includes transcriptional regulators that define early endodermal and epithelial programs, such as *Sox17* [47], *Foxa2, Gata4, Hnf1b*, and *Hnf4a*, which are central components of early endodermal patterning and epithelial specification networks [48–50]. The cluster also contains genes associated with epithelial function and metabolic organization, including *Fabp1, Muc13, Aqp8*, and *Enpep*,reflecting functional programs that accompany early epithelial differentiation [51].

The persistence of localization across early time points suggests stable organization of these transcriptional and functional programs during initial patterning, whereas the loss of localization by day 10 indicates redistribution of these programs as later, more specialized cell states emerge [52].

Together, these temporal localization trajectories provide a dynamic view of cellular differentiation as a complex reorganization where early transcriptional programs dissolve while new lineage-specific organization emerges, marking distinct developmental transitions across time, which we detect as localization events on embedding of intermediate time-points.

#### Gene localization analysis highlights a developmental watershed between day 4 and day 10

The most pronounced reorganization of gene localization occurs between day 4 and day 10. Although comparable numbers of genes were localized at each stage (1323 at day 4, and 1429 at day 10), only 529 genes were shared between these timepoints, indicating a substantial restructuring of transcriptional organization. This shift reflects a major transition in cellular state organization during retinoic acid–driven differentiation. As described above, genes enriched in day 4 over-represented clusters (clusters 0010 and 1110) are associated with early patterning, epithelial, and progenitor-associated programs, whereas genes localized specifically at day 10 (cluster 0001) reflect increasingly organized neural lineage programs. The near-complete turnover of localized gene sets suggests that the interval between day 4 and day 10 marks a developmental watershed characterized by the resolution of early, multi-lineage transcriptional organization and the emergence of more lineage-restricted neural programs. This transition is consistent with established retinoic acid differentiation dynamics, in which early patterning and endoderm-associated programs diminish as neural identities become increasingly dominant [53].

#### Differential localization enables tracking of stable cell populations across developmental transitions

Beyond identifying stage-specific programs, temporal localization analysis reveals a particularly informative form of differential localization: genes that are expressed across conditions but transition from non-localized to localized as cellular heterogeneity increases. This pattern is exemplified by *Id3* and *Cldn3*, which exhibit distinct but complementary localization trajectories over the differentiation time course (Fig. 5). *Id3*, a regulator associated with proliferative progenitor states [54], is broadly expressed and weakly or non-localized at early time points but becomes localized as progenitor populations begin to segregate during differentiation. In contrast, *Cldn3* [55], which encodes a tight-junction component linked to epithelial organization, is unlocalized at day 0, becomes weakly structured at intermediate stages, and is strongly localized by day 10.

These changes do not imply altered gene function or cell-intrinsic regulation. Rather, they reflect shifts in *population context*. At early time points, the culture is dominated by relatively homogeneous, cycling pluripotent cells, causing broadly required regulators and structural components to be widely expressed [56]. As differentiation proceeds, increasing cellular diversity and the emergence of post-mitotic lineages restrict both proliferative and epithelial programs to specific subpopulations, allowing genes such as *Id3* and *Cldn3* to exhibit localized expression patterns [57–59].

The ESC differentiation dataset provides a particularly clear example of this regime of differential localization. As differentiation proceeds and population composition diversifies, genes that are broadly associated with early progenitor populations become increasingly confined to specific subpopulations. Localization often emerges gradually: genes may exhibit modest confinement at intermediate time points before becoming more strongly localized as additional cell states arise. With increasing heterogeneity, these genes are not only enriched within particular lineages but also depleted from surrounding states. Detecting such transitions therefore requires evaluating both concentration within a target population and absence elsewhere.

This behavior is illustrated by genes such as *Id3* and *Cldn3*, which show progressive confinement to progenitor or epithelial subpopulations across the time course.Although their degree of localization varies across stages, their enrichment within specific lineages is accompanied by reduced expression in emerging states. By jointly capturing enrichment and depletion, localization analysis enables identification of cell populations that persist across developmental transitions despite substantial shifts in overall population composition.

## 3 Discussion

In the present work we refine the definition of a “marker gene” in single-cell analysis as a gene whose expression is both concentrated within a specific subset of cells and significantly depleted in complementary subsets. This refinement addresses a limitation of enrichment-based approaches, which can identify genes that vary across cells yet remain broadly expressed across much of the population. Such genes may exhibit gradual trends or diffuse upregulation without clearly delineating distinct cell populations. By explicitly requiring statistical evidence of depletion outside enriched neighborhoods, we prioritize genes whose expression is restricted to well-defined subsets rather than broadly distributed. In this framework, depletion provides the statistical criterion that distinguishes specific population markers from genes that merely reflect structured variation.

Existing marker gene selection methods use a variety of approaches: gene autocorrelation in embedding space (Hotspot [3]), density-based scoring (singleCell-Haystack [2]), differential expression tests (Scanpy [9]), and diffusion-based metrics (LMD [5], GSPA [6]). Yet, with the exception of LMD and GSPA, which implicitly penalize background expression through diffusion- or pattern-based scoring (Section 2.4), most of these methods are designed around enrichment-oriented signals rather than an explicit combination of enrichment and depletion evidence. Our approach, Locat, tests both concentration and depletion simultaneously through density contrasts, distinguishing genes that satisfy this dual criterion from genes that concentrate locally but retain substantial background expression.

Locat identifies localized genes whose expression exhibits sharp contrast against the global background density. These genes therefore form a highly specific, compact, and interpretable feature set within each sample that is well suited for visualization and downstream analyses. In multi-sample studies, we find it most informative to perform marker gene discovery separately within each sample’s embedding, then compare localization scores across samples and, when appropriate, construct joint visualizations using the union of condition-specific localized genes. This strategy avoids applying batch correction during marker gene discovery and directly addresses a central challenge in current analytical pipelines: joint embeddings without batch correction are often dominated by sample-specific and technical variation, while batch-correction–based integration can suppress or obscure the condition-specific gene programs that experimental designs aim to study.

Where controlled perturbations are applied to shared lineages, comparing localization scores across independently embedded samples offers a conservative alternative to aggressive integration. For multi-sample datasets with higher variation between samples, or time courses with varying cell population composition, we demonstrate that identifying localized genes in each sample separately enables identification of similar populations across samples. This enables researchers to compare samples through the lens of marker genes in order to e.g. track emergent cell populations or to link minority subpopulations from later developmental stages to their ancestral proliferating cells.

Locat inherits limitations from both density modeling and the embedding geometry on which it operates. Inference is performed in the user’s chosen coordinate system, and distortions of distance scales or neighborhood relationships can therefore influence both concentration and depletion evidence. In this work, we primarily fit models in higher-dimensional PCA coordinates and used UMAP mainly for visualization, as low-dimensional nonlinear embeddings may warp global distances and relative scales in ways that are not faithful to the data. We therefore recommend fitting Locat on embeddings that are well supported for the biological signals of interest, prioritizing representations with stable geometry, and interpreting conclusions that depend on fine-grained distance relationships in two-dimensional nonlinear embeddings with caution. At the same time, cross-fold reproducibility analyses in the IFN-*β* dataset indicate that localization rankings are reasonably stable across the tested embedding choices, including 4, 8, and 12 PCA dimensions and a BiPCA →UMAP setting, with cross-fold AUROC values of 0.923–0.979 and top-500 Spearman correlations of 0.681– 0.809 (Supplementary Tables S3 and S5), while within-fold seed stability remained similarly high across embedding choices (Supplementary Table S4). Another practical consideration is expression prevalence: genes expressed in very few cells provide limited effective sample sizes, which can reduce statistical power and increase sensitivity to modeling choices. While our simulation-based calibration explicitly accounts for the dependence of localization statistics on expression prevalence and includes effective sample size diagnostics, signals from extremely rare genes remain challenging for any localization-based approach.

Several directions for future work arise from this framework. The concentration–depletion logic may be adapted to spatial transcriptomics data, where physical tissue coordinates provide a natural representation for assessing localized expression. In such settings, testing for enrichment within spatial neighborhoods and depletion outside them could help distinguish genes marking anatomically confined regions from those that are more broadly distributed across tissue. Related opportunities exist in multi-modal assays such as single-cell multiome or CITE-seq, where evaluating whether transcriptional localization is accompanied by coordinated restriction in chromatin accessibility or protein abundance may provide stronger evidence for coherent regulatory programs. Finally, extending the framework to evaluate sets of genes jointly, rather than testing each gene independently, may enable identification of module-level markers defined by coordinated expression patterns. Together, these possibilities suggest that the concentration–depletion principle may provide a unifying perspective for identifying specific biological programs across diverse single-cell modalities.

## 4 Materials and Methods

### 4.1 Locat

#### 4.1.1 Input Data

Locat takes two inputs: a gene by cell normalized count matrix (the expression matrix) *W* ∈ ℝ^*m×n*^, where *G* is the set of all genes (|*G*| = *m*) and *C* is the set of all cells (|*C*| = *n*), and a *d*-dimensional cell embedding *X* ∈ ℝ^*n×d*^, such that each cell *c* ∈ *C* has normalized count vector *w*_*\*,c*_ and embedding coordinate **x**_*c*_ ∈ ℝ^*d*^. Locat evaluates localization with respect to the embedding *X* using normalized counts *W* as informative weights. Internally, Locat standardizes each embedding dimension to zero mean and unit variance before fitting the background and gene-specific mixture models, so that density estimation is not dominated by dimensions with larger raw scale.

#### 4.1.2 Density estimation

First, Locat models both the background cell density and gene-specific expression density using weighted Gaussian mixture models (WGMMs, Supplementary Methods S1.1). By default, the background WGMM uses uniform per-cell weights; optionally, sample-specific weights (for example, total expression count per cell) can be used. Gene-specific WGMMs use the expression level of that gene across cells. In all cases, weights are normalized to sum to one. We use BIC (Supplementary Methods S1.1.3) as a model-selection criterion that balances goodness of fit and model complexity for selecting background model complexity and guiding component selection.

The background WGMM *f*_0_(**x**) is fit once per dataset using all cells in the embedding. We average over multiple random initializations to stabilize the estimated background density values *f*_0_(**x**_*c*_) at each cell *c*. For each gene *g*, we fit a signal WGMM *f*_*g*_(**x**) using per-cell weights *w*_*g,\**_ normalized to sum to one. The number of components in this gene-specific model is adapted to the expression pattern of each gene: genes spread across larger regions of the embedding are assigned more components,while compact patterns use fewer components (Supplementary Methods S1.1.4).

#### 4.1.3 Concentration Score

Next, we quantify the concentration score, which evaluates whether gene expression density concentrates into localized peaks within the embedding. We first compute a normalized difference between background and signal density. We then convert this raw score to a z-score by comparing it to a null distribution built from randomized pseudogenes with similar prevalence (fraction of expressing cells), so genes are compared fairly across different detection rates. Finally, we compute statistical significance from that z-score. Before scoring, genes with prevalence above a user-defined upper bound are excluded (parameter `max freq,` default 0.9).

Let *ϕ*_*g*_ be the fraction of cells with *w*_*gc*_ *>* 0, which is the expression prevalence of gene *g*. Recall that *f*_0_(**x**_*c*_) is the background density at cell *c*, and *f*_*g*_(**x**_*c*_) is the gene-specific density for gene *g* at cell *c*.

Concentration is based on a density contrast score comparing gene-specific and background densities, defined as:

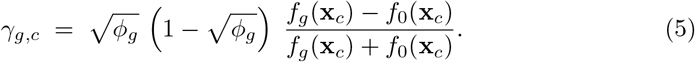

This quantity is positive where the gene density exceeds the background and negative otherwise. The prevalence factor 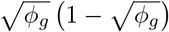 stabilizes the statistic for sparse genes while downweighting broadly prevalent genes.

The per gene concentration statistic is the expression weighted average of *γ*_*g,c*_ over expressing cells,

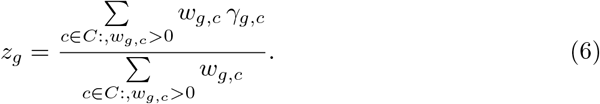

We then adjust *z*_*g*_ to account for prevalence as described in Supplementary Methods S1.2. Specifically, we estimate *µ*(*ϕ*) and *σ*(*ϕ*) from randomized pseudo-genes generated across prevalence levels and interpolate these functions over *ϕ*. For gene *g* with prevalence *ϕ*_*g*_, the prevalence-adjusted concentration z-score is:

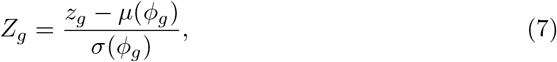

and compute the one-sided concentration *p*-value

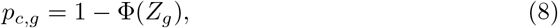

where Φ is the standard normal cumulative distribution function. Genes whose expressing cells lie in regions where *f*_*g*_ is consistently larger than *f*_0_ result in a large positive *Z*_*g*_ and therefore small *p*_*c,g*_.

#### 4.1.4 Depletion Score

The second term in our dual criterion is the depletion score. It tests whether a gene is under-represented in parts of the embedding where many cells are expected under the background model. We evaluate this at several contrast levels *λ* ≥ 1 to examine different region definitions: lower *λ* values include larger regions of depletion that may extend toward boundaries between depleted and enriched areas, whereas higher *λ* values define smaller, stricter regions that are more clearly separated from regions of enrichment. For each *λ*, we define a region of depletion and compute two fractions in that region: (i) the expected fraction under the background model and (ii) the observed fraction among cells expressing the gene. Using these quantities, we perform a one-sided depletion test for that contrast, where the statistic is more significant when the observed fraction is further below the expected fraction under the null model. The corresponding *p*-value uses an effective sample size adjustment to account for unequal weights and dependence among nearby cells in the embedding, which would otherwise underestimate variance and inflate significance. Because multiple *λ* values are evaluated for each gene, we apply a multiple-testing correction across that scan.

##### Regions of relative depletion and expected density mass

Let *f*_0_ and *f*_*g*_ be the background and gene *g* WGMM density functions respectively. Given a specific threshold *λ* ≥ 1, termed the contrast threshold, we determine the *region of depletion* as the set of cells Γ_*λ*_ for which the background density exceeds the gene-specific density by a factor at least equal to *λ*:

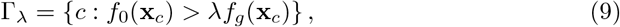

The expected background mass represents the fraction of total background density falling within the region of depletion:

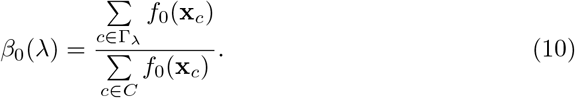

The observed gene-specific mass for gene *g* is the fraction of expressing cells falling within the region of depletion:

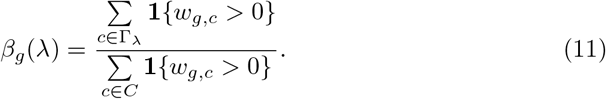

The two definitions above are intentionally different. *β*_0_(*λ*) is model-based: it is computed from fitted background density values *f*_0_(**x**_*c*_) and represents the expected fraction of background density contained in Γ_*λ*_ under the null. *β*_*g*_(*λ*) is empirical: it is computed from observed expressing cells via the indicator **1 {***w*_*g,c*_ *>* 0 }and represents the observed fraction of expressing cells in the same region of depletion. Under the null, these two quantities should be similar. Evidence for depletion is obtained when *β*_*g*_(*λ*) is substantially smaller than *β*_0_(*λ*). In the default analysis, each expressing cell contributes equally to *β*_*g*_(*λ*), so depletion is tested on expression frequency rather than expression magnitude. Optionally, cells can instead be weighted by nonnegative expression values, yielding a depletion test on total expression amount.

Both expected and observed quantities are computed by summing over the observed cells in the dataset, rather than integrating over the full continuous embedding space. This keeps the test tied directly to the cells and embedding positions that were actually measured. Integrating over the full embedding would be computationally expensive and would assign weight to regions where no cells were observed.

##### Tail probability

Under the null hypothesis that cells expressing gene *g* are distributed in proportion to the background density, we compute a one-sided lower-tail probability using a beta-binomial model with effective trial count *n*_trials_ = round(*n*_eff_). For each contrast *λ*, let *k*_obs_(*λ*) = round(*β*_*g*_(*λ*) *n*_trials_) and *p*_0_(*λ*) = *β*_0_(*λ*). Here *ρ*_bb_ is a fixed overdispersion parameter that controls extra variability relative to a binomial model; the default setting is *ρ*_bb_ = 0.02. We define

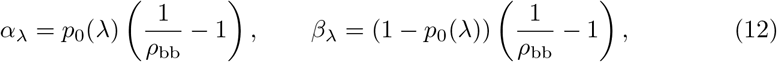

and evaluate

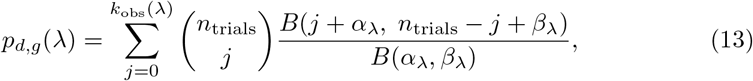

where *B*(·, ·) is the beta function. If *ρ*_bb_ = 0, the test uses a binomial lower-tail calculation. Here *n*_eff_ is Kish’s effective sample size for the given gene *g*,

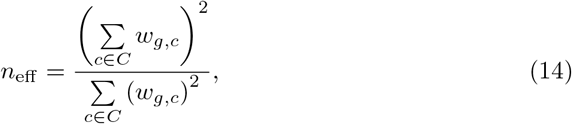

with *w*_*g,c*_ being the normalized count vector for gene *g* in cell *c*. Equation 14 reflects the reduction in effective degrees of freedom when observations carry unequal weights. Using the raw cell count *n* would treat all cells as equally informative and yield overly permissive *p*-values when a small number of highly weighted cells dominate the test statistic. To further ensure stability and conservativeness, we apply a multiplicative scaling factor to *n*_eff_ and then impose an upper bound on the scaled *n*_eff_](see Supplementary Methods for parameterization details). In the primary analyses reported here, we used a fixed effective-sample-size scaling factor of 0.6, an upper cap of 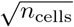 son effective trials, and a beta-binomial overdispersion parameter *ρ*_bb_ = 0.02

(Supplementary Table S1). These settings were held constant across datasets as fixed analysis settings rather than tuned per analysis, and their combined effect is assessed empirically through the null-calibration analyses in Supplementary Fig. S1. These adjustments account for the fact that *f*_0_ and *f*_*g*_ are not independent: *f*_*g*_ is fit using a subset of the same cells that define *f*_0_, and both densities are fit to cellular positions in the embedding, where nearby cells share expression patterns.

##### Selecting a contrast

Genes may exhibit significant depletion at different contrast thresholds, depending on the shape of their density profiles and how sharply expression declines outside concentrated regions. We therefore evaluate the depletion tail probability over a linearly spaced grid of *λ* values with *λ* ≥ 1, using a moderate finite range of contrasts, and account for this scan in the final p-value calculation. To ensure that the test is well-conditioned and supported by sufficient evidence, we restrict attention to the subset Λ of *λ* values satisfying the following criteria:

- the background density mass *β*_0_(*λ*) in the region is sufficiently large to support a meaningful comparison;
- the expected effective count *n*_eff_ *β*_0_(*λ*) exceeds a minimum sample-size threshold;
- the absolute deficit *β*_0_(*λ*) −*β*_*g*_(*λ*) exceeds a minimum threshold;
- the observed gene-specific mass does not exceed the level implied by the contrast ratio 1*/λ*, i.e., *β*_*g*_(*λ*) *<* (*β*_0_(*λ*)*/λ*) (1 − *ε*_rel_).

These conditions exclude contrasts that produce extremely small regions, unstable effective sample sizes, or negligible deviations from the null.

Among admissible contrasts, we retain the minimum *p*_*d,g*_(*λ*) over Λ. If no contrast satisfies the above conditions, the depletion p-value is set to 1.

To adjust for multiplicity across scanned *λ* values, we apply a Šidák-form adjustment based on the effective number of tested contrast thresholds *n*_Λ_, which results in the depletion score:

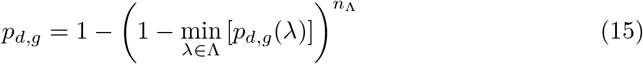

Because tests across *λ* are correlated, this adjustment should be interpreted as an approximate multiplicity correction across tested *λ* values. Small *p*_*d,g*_ indicates that gene *g* is under-represented across a substantial portion of the background density (high *f*_0_), consistent with exclusion from areas outside its concentrated regions.

#### 4.1.5 Localization Score

Finally, we combine the concentration and depletion scores to obtain one localization score. The localization score is adjusted with penalization terms aimed at improving robustness for small sample-sizes.

In detail, a raw localization score is computed using a Cauchy combination test [60], which is meant to produce a valid omnibus *p*-value even under complex dependence:

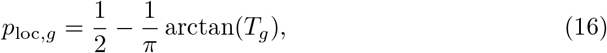

where

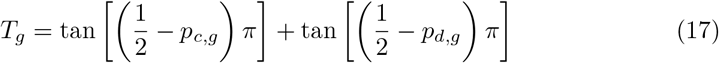

To compute the adjusted localization score we incorporate two small penalty terms that stabilize the combined statistic. The first penalizes genes supported by very few expressing cells, for which density estimates can be unstable. The second penalizes genes whose fitted signal density does not dominate the background at most of their expressing cells, thereby discouraging diffuse or poorly separated expression patterns even within their putative regions of concentration.

Let *n*_*g*_ =Σ _*c∈C*_ **1** {*w*_*g,c*_ *>* 0} be the number of cells expressing gene *g* i.e., the support of gene *g*. We define the sparsity adjustment factor

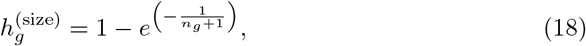

which is close to 0 for well sampled genes and larger for rare genes. We also define the sensitivity adjustment factor

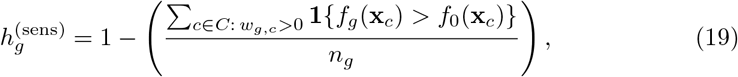

as the fraction of cells expressing gene *g* for which the gene density exceeds the background density.

The adjusted localization score i.e., the output from Locat, is

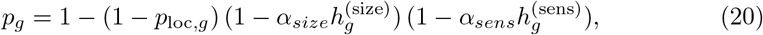

clipped in (0, 1). The coefficients *α*_size_ and *α*_sens_ control the strength of sparsity and sensitivity penalization, respectively; in all analyses reported here we used *α*_size_ = 0.05 and *α*_sens_ = 0.12. Default parameter values and robustness analyses are described in Supplementary Methods S1.4 and Supplementary Fig. S22. Finally, we apply a monotone smoothing map near the upper tail:

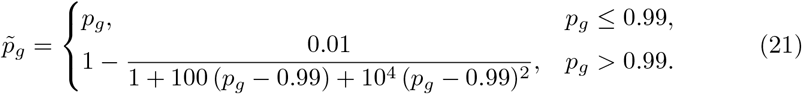

This transformation only affects values very close to one and preserves rank ordering, while preventing genes from collapsing to nearly identical upper-tail values. As a result, it primarily improves numerical separation among insignificant genes and has negligible effect on which genes are called significantly localized. The reported final localization *p*-value is 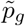.

By construction, *p*_*g*_ is a conservative adjusted localization score: Eq. 20 is monotone and satisfies *p*_*g*_ ≥ *p*_loc,*g*_. Thus, the penalty terms can only increase the score relative to the Cauchy-combined value, stabilizing rankings for sparse or weakly separated genes rather than introducing additional significance. Robustness of gene rankings to the penalty coefficients *α*_size_ and *α*_sens_ is shown in Supplementary Methods S1.4 and Supplementary Fig. S22.

#### 4.1.6 Locat Output

For each gene *g*, Locat is designed to explicitly report:

- *p*_*g*_: adjusted localization score (eq. 20);
- *p*_*c,g*_: concentration significance (eq. 8);
- *p*_*d,g*_: depletion based localization significance (eq. 15);
- *Z*_*g*_: prevalence-adjusted concentration *z*-score (eq. 7);
- a BIC-like score that quantifies the parsimony of the fitted WGMM (Supplementary Methods S1.1.3);
- *K*_*g*_: estimated number of mixture components in the gene’s signal WGMM (Supplementary Methods S1.1.4);

Genes with *p*_*g*_ below a chosen significance threshold are considered localized. In this setting, we evaluate calibration empirically under simulation-based null analyses (Supplementary Methods S1.3) rather than applying a single genome-wide correction across genes. This is because the gene-level tests are not an exchangeable family: all genes are tested in the same embedding with a shared background/gene-specific GMM density framework, while prevalence and noise vary substantially across genes.

The adjusted localization score 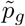 combines the concentration and depletion p-values into a single monotone score, using two penalty coefficients chosen once from preliminary simulations rather than tuned per dataset. It should therefore be read as a ranking and thresholding guide rather than a *p*-value with a formally proven error rate. Simulation-based null analyses matched to Locat’s pipeline structure support this use, showing that the 0.05 threshold is reasonably well calibrated across the null scenarios tested (Supplementary Methods S1.3, Supplementary Fig. S1).

### 4.2 Modules identification

To identify clusters of co-expressed genes, we extract the gene-by-cell expression matrix for a specified set of input genes. We binarize the expression matrix using a threshold on expression levels. For each gene pair, we quantify overlap between the resulting binary expression vectors across cells and evaluate whether the observed co-expression across cells exceeds baseline expectation using a hypergeometric test. We retain gene pairs with significant co-occurrence across cells as positive edges to form a binary gene–gene adjacency matrix. Louvain community detection then partitions this graph to recover gene modules i.e., clusters of genes with substantial co-expression.

In detail, the entries *b*_*g,c*_ of the binarized expression matrix *B* = (*b*_*g,c*_), derived from the normalized expression matrix *W* as follows:

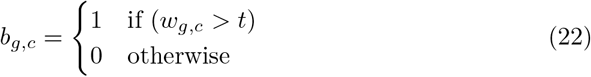

where *G* is a pre-defined list of genes, *w*_*g,c*_ is the expression level for gene *g* in cell *c*, and *t* is a threshold on gene expression. In our analyses *t* = 0.

The entries 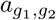 of the gene-gene adjacency matrix, where *g*_1_ and *g*_2_ are two genes, are:

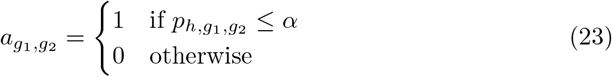

where 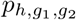 is the hypergeometric p-value for the overlap between binary expression vectors 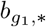 and 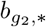, and *α* is the chosen significance threshold, in our analyses *α* = 0.05.

To ensure interpretability and stability, communities containing very few genes are discarded. The remaining communities represent gene modules.

### 4.3 Data Preprocessing

Datasets were accessed as annotated AnnData or Seurat objects when available. When preprocessed matrices or embeddings were not provided, we applied standard normalization and dimensionality reduction to obtain inputs for Locat. In all analyses, we used the log-normalized count matrix as the source of cellular weights in the gene-specific WGMM models. Prior to analysis, we removed genes expressed in fewer than 1% of cells and excluded genes with symbols indicative of predicted, uncharacterized, or non–protein-coding loci (e.g., Gm-, Rik-, AC-/AA-prefixed genes and related annotations), retaining a predominantly protein-coding gene set. For embedding construction, we used the first several principal components derived from the log-normalized expression matrix, typically between 4 and 8 components across datasets.

### 4.4 Neighborhood Differential Abundance

To identify neighborhoods associated with condition- or cohort-specific enrichment, we performed neighborhood differential abundance (nDA) analysis, a neighborhood overrepresentation procedure adapted from Landa *et al*. [61]. Given cell coordinates *X* ∈ ℝ^*n×d*^ and a binary (or probabilistic) group assignment, nDA produces a final per-cell score 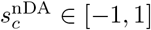, where 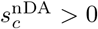 indicates local enrichment and 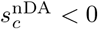 indicates local depletion of the group of interest.

Let *y*_*c*_ ∈ {0, 1} denote the hard group label for cell *c* (or *q*_*c*_ ∈ [0, 1] a soft membership probability). We first build a single `NearestNeighbors` index on *X* and query *k*-nearest-neighbor sets *N* _*k*_(*c*) for *k* ∈ {15, 40, 75 }. For each cell *c* and each scale *k*, we compute the local group-1 fraction

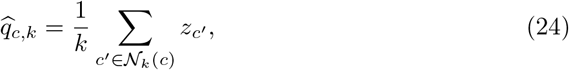

where 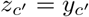for hard labels and 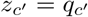for soft labels. We also compute the global group-1 prevalence

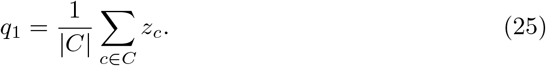

We convert each (*c, k*) into a signed enrichment score using a Bayes-rule-derived transform,

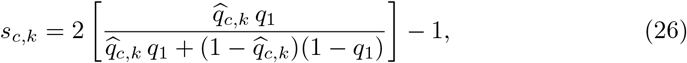

yielding a multiscale score matrix **S** = [*s*_*c,k*_].

Scores are computed for each *k* in parallel chunks to reduce memory overhead. To aggregate information across scales, we fit a logistic regression classifier to predict group membership from **S**, and linearly rescale the fitted posterior probabilities to [− 1, 1] to obtain the final nDA score 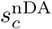 per cell. This multiscale logistic aggregation increases sensitivity to neighborhood-size heterogeneity while preserving the direction of enrichment.

Implementation details: nDA was implemented in Python using `scikit-learn` for nearest-neighbor search and logistic regression. Analyses were performed on PCA coordinates with Euclidean distance, using balanced class weights. In all experiments, we evaluated neighborhood sizes *k* ∈ {15, 40, 75 } and used default logistic regression settings.

### 4.5 Embeddings Derived from Gene Subsets

In selected analyses, we constructed additional low-dimensional embeddings restricted to specific gene subsets, such as localized genes identified by Locat or predefined gene partitions (e.g., differentiation trajectory genes in embryonic dermis; see Section 2.6). To do so, we subset the log-normalized expression matrix to the genes of interest, recomputed principal components from this restricted matrix, and generated two- or three-dimensional UMAP embeddings for visualization. When multiple experimental conditions were analyzed jointly (e.g., control and stimulated samples in interferon-*β* stimulation data; see Section 2.7), embeddings were constructed from the combined dataset using the shared gene subset. These subset-derived embeddings were used for visualization and comparative analysis.

### 4.6 Likelihood-ratio Test for localization

As a complementary diagnostic for gene localization, we implemented a likelihood-ratio test (LRT) comparing the gene-specific weighted Gaussian mixture model (WGMM) density to the background WGMM density on the full embedding (see Supplementary Methods S1.1). Both models are fit as described above, using the same embedding coordinates and covariance regularization strategy described in Section 4.1.2.

For a given gene *g*, let ℐ_*g*_ = {*c* : *w*_*g,c*_ *>* 0 }denote the set of cells expressing the gene. We evaluated the log-likelihood of these cells under the background model, *f*_0_(**x**_*c*_), and under the gene-specific signal model, *f*_*g*_(**x**_*c*_). The likelihood-ratio statistic was computed as

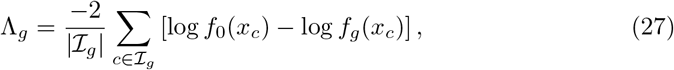

where normalization by |ℐ_*g*_| ensures comparability across genes with different expression prevalences.

Under the null hypothesis that the distribution of expressing cells in the embedding is consistent with the global background structure, Λ_*g*_ was compared to a chi-square distribution with degrees of freedom

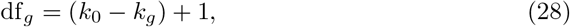

where *k*_0_ and *k*_*g*_ denote the number of mixture components in the background and gene-specific models, respectively. The corresponding likelihood-ratio *p*-value is

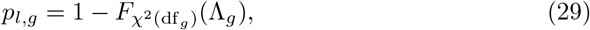

where 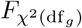 is the chi-square cumulative distribution function with df_*g*_ degrees of freedom.

Genes expressed in fewer than a minimum number of cells or in more than a specified fraction of cells were excluded from testing. In practice, the likelihood-ratio test was used to quantify the extent to which a gene’s expressing cells deviate from the background embedding distribution, but was not used as the primary criterion for localization in Locat.

### 4.7 Permutation Test for localization

Although local and global gene scores are generally conservative in the standard Locat model, we also provide permutation-based empirical p-values as an optional diagnostic filter. These methods estimate localization *p*-values for each tested gene and can be computationally expensive, so in practice we suggest applying them to a limited set of candidate genes that already pass the combined localization threshold in the standard Locat model. For each null replicate, we sort cells by the gene’s normalized weights, divide them into rank-based bins of similar weight magnitude, and permute weights among cells within each bin. We then multiply these shuffled weights by an independent positive random factor per cell, introducing bootstrap-like variability in each cell’s effective contribution, and refit the WGMM for that replicate. Empirical *p*-values are computed by comparing the observed localization statistic to the replicate-based null distribution across *n*_*initializations*_ runs (100 by default).

### 4.8 Benchmarking workflow

We consider multiple clustering-free tools as well as Locat in several benchmarking experiments. Unless otherwise specified, each of these tools uses the same AnnData object [9], input gene list, and log-normalized reads as input, and uses nearest neighbors, PCs, and/or UMAP coordinates derived from these data. We therefore keep relatively consistent preprocessing steps to ensure each method is fairly evaluated. For ranking comparisons we always sort by the most significant corrected *p*-value/FDR for each method.

#### Scanpy-DE

Louvain is applied to the neighborhood graph using default parameters in Scanpy [9], yielding a dictionary of clusters assignments. For each cluster, we use default parameters to calculate one-vs-all differential expression tests the Wilcoxon rank-sum Differential Expression tests over all genes. We then rank genes by significance, using the most significant corrected *p*-value for each gene (thus, if a gene is significant in two clusters and not significant in any others, the smallest of the two significant *p*-values will be used to rank the gene against other evaluated genes).

#### singleCellHaystack

We used default parameters in the Python implementation of singleCellHaystack, using PCs as the input embedding parameter in the *train haystack logpv* function. We then rank the genes by sorting the *log pvalue* output.

#### Hotspot

We used the Python implementation of Hotspot, specifying the “normal” model be used to measure gene autocorrelation. We then recompute the KNN graph and compute gene autocorrelations. In order to rank the genes, we used the computed false discovery rate (FDR) as a proxy for significance of a given gene.

#### GSPA

We used the Python implementation of GSPA, following the GitHub repository and examples as a guideline for parameter selection. We compute the PHATE operator [62] as the input embedding to GSPA, similarly to the examples on the GitHub repository. We then use the *gspa*.*GSPA*.*calculate localization*() function to return localization probabilities per each gene. We found that GSPA was sometimes unstable when trained with large data objects. We created a pseudo-GSPA method that enabled sparse input to the *gspa*.*GSPA*.*construct graph*() method and allowed us to calculate *gspa*.*GSPA*.*get gene embeddings*() in a sparse, batched format.

#### LMD

We used the published R implementation of LocalizedMarkerDetector (LMD), following the GitHub repository and tutorial workflow as a guideline for inputs and parameter selection [5]. We passed LMD a genes-by-cells expression matrix and a cells-by-dimensions PCA feature space, and used the default kNN-based graph construction to compute per-gene cumulative localization scores returned by *LocalizedMarkerDetector* :: *LMD*. To ensure compatibility when calling LMD from Python, we implemented an rpy2 wrapper that constructs R matrices with unique gene and cell identifiers, filters genes expressed to a chosen minimum number of cells, and removes ultra low-variance principal components that can otherwise cause failures. For large datasets we additionally used LMD’s coarse-grained large-data workflow (*CoarseGrain* and *fast get lmds*) to compute the same cumulative scores efficiently.

#### GiniClust

We used the Python implementation of GiniClust3 [63] to compute the Gini coefficient of each gene’s expression across cells (*giniclust*3.*gini*.*giniIndex*) on the same log-normalized expression matrix used by the other methods. Genes were then ranked by decreasing Gini coefficient, such that genes with more unequal, concentrated expression across cells are ranked as more specific.

#### SpectralRH

We implemented SpectralRH as a baseline geometric metric defined in this work (see Supplementary Methods S1.8). For each gene, we calculated Rayleigh smoothness and spectral entropy on the cell–cell graph built from PCA coordinates using Scanpy’s `pp.neighbors()` function to construct the normalized Laplacian. We used the first *m* = 96 Laplacian eigenvectors (a fixed default applied uniformly across all datasets in this comparison, not tuned per dataset) to compute spectral entropy and combined the two measures after sample-size–matched null calibration to obtain a single SpectralRH score per gene. Genes were then ranked by increasing SpectralRH value, such that lower scores correspond to smoother, more locally coherent, and potentially localized expression patterns.

#### Cell-type specificity score

To evaluate the cell-type specificity of gene sets returned by each method, we define the cell-type specificity score *S*_*g*_ for gene *g* as

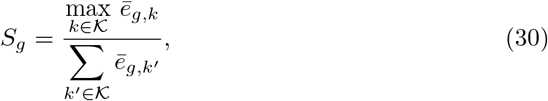

where *K* is the set of annotated cell types and 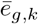 is the mean log-normalized expression of gene *g* in cell type *k. S*_*g*_ is defined for genes with positive total expression and ranges from 0 to 1, with values near 1 indicating expression almost entirely confined to a single cell type. Genes expressed in fewer than a dataset-specific minimum fraction of cells are excluded. We used a 5% expression threshold for PBMC3k and dermal condensates, and retained all genes passing the *n*_cells_ ≥ 100 filter for the Kang datasets. For the *Chlamydomonas* Fe^+^ and Fe^*−*^ datasets and the Tabula Muris Senis bone marrow dataset, a 5% expression threshold was used, with cell-type groupings defined by Leiden clustering (resolution 0.3) and annotated cell types, respectively.

#### 4.8.1 Empirical significance calculations

Neither GSPA nor LMD directly return *p*-values for gene-level significance. To enable direct comparison across methods and datasets, we converted raw localization scores into empirical significance values using within-sample gene rankings. For each method, genes were ranked by their reported score, with more extreme values indicating stronger evidence for non-uniform expression. Empirical *p*-values were computed as

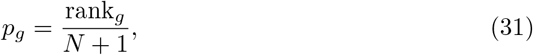

where *N* is the total number of genes evaluated and ranks were computed using average ranking to handle ties. For LMD, lower cumulative scores correspond to stronger signal, and ranks were therefore computed in ascending order. This rank-based calibration preserves each method’s internal gene ordering while providing a standardized scale for cross-method comparison.

## Supporting information

Supplementary Info including data, figures, methods.

## 4.9 Data availability

All datasets analyzed in this study are publicly available from their original publications. The murine embryonic dermis dataset (GEO: GSE255534) was obtained from [7, 16]. The IFN-*β* stimulation PBMC dataset (GEO: GSE96583) was obtained from [17]. The retinoic acid differentiation time course (GEO: GSE227320) was obtained from [28]. The 10X PBMC 3k dataset was obtained from [8]. The *Chlamydomonas reinhardtii* Fe^+^/Fe^*−*^ transcriptome (GEO: GSE157580) was obtained from [10]. The Tabula Muris Senis bone marrow dataset was obtained from [11] via the CZI CELLxGENE Census.

## 4.10 Code availability

Code for running Locat is available via PyPI at https://pypi.org/project/locat and https://github.com/KlugerLab/Locat, and notebooks for generating results in this article are available at https://github.com/KlugerLab/Locat-paper. Analyses reported here were run against `locat` v0.1.5.

## 4.11 Acknowledgements

The authors thank Erez Peterfreund, Peggy Myung, and Boaz Nadler for valuable discussions on related data analysis and methodology, and for insightful feedback on the development of the localization framework.

