## Supplementary Info including data, figures, methods. for "Locat: Joint enrichment and depletion testing identifies localized marker genes in single-cell transcriptomics"

### Supplementary Information

**S2 Supplementary Data** **9**

**S3 Supplementary Figures** **10**

**S4 Supplementary Tables** **28**

**S5 Supplementary Notes** **32**

### S1 Supplementary Methods

#### S1.1 Weighted Gaussian Mixture Models (WGMMs)

A Gaussian mixture model (GMM) assumes observations are generated from a finite mixture of Gaussian distributions with unknown parameters. A weighted GMM (WGMM) extends this model by incorporating sample weights that encode the relative importance of observations.

##### S1.1.1 Model.

The probability density function of a $K$-component WGMM is a weighted sum of $K$ Gaussian component densities:

$$f(\mathbf{x}) = \sum_{k=1}^{K} \pi_k \, \mathcal{N}(\mathbf{x} \mid \mu_k, \Sigma_k), \tag{1}$$

where $\mathbf{x}$ is a point in sample space and $K$ is the number of components. Here, $\pi_k \geq 0$ is the mixture weight of component $k$ with $\sum_{k=1}^{K} \pi_k = 1$, and $\mu_k$ and $\Sigma_k$ are that component's mean and covariance, respectively. In the main Methods, the background density $f_0(\mathbf{x})$ and gene-specific density $f_g(\mathbf{x})$ are both instantiated as WGMMs under this formulation.

**Model estimation via weighted EM updates.** Model fitting requires estimation of $(\pi_k, \mu_k, \Sigma_k)$, performed iteratively using the Expectation-Maximization (EM) algorithm.

In the notation below, we omit iteration indices; the symbol $\leftarrow$ denotes parameter updates.

Given parameter estimates at iteration $t$, $(\pi_k, \mu_k, \Sigma_k)$, the E step computes responsibilities:

$$r_{ik} = \frac{\pi_k \, \mathcal{N}(\mathbf{x}_i \mid \mu_k, \Sigma_k)}{\sum\limits_{\ell=1}^{K} \pi_\ell \, \mathcal{N}(\mathbf{x}_i \mid \mu_\ell, \Sigma_\ell)}, \tag{2}$$

which are the posterior probabilities that cell $i$ belongs to component $k$.

Sample weights $s_i$ enter the M step through effective weighted responsibilities:

$$v_{ik} = s_i \, r_{ik}, \tag{3}$$

such that higher weighted cells exert proportionally stronger influence.

Let $w_k = \sum_i v_{ik}$ and $u = \sum_{k=1}^{K} w_k$. The mixture parameters are then updated as

$$\pi_k \leftarrow \frac{w_k}{u}, \tag{4}$$

$$\mu_k \leftarrow \frac{\sum_i v_{ik} \mathbf{x}_i}{w_k}, \tag{5}$$

$$\Sigma_k \leftarrow \frac{\sum_i v_{ik}(\mathbf{x}_i - \mu_k)(\mathbf{x}_i - \mu_k)^\top}{w_k} + \lambda I, \tag{6}$$

where $\lambda$ is a small regularization term to ensure numerical stability. These updates match the standard weighted-EM formulation, and our implementation follows Gebru et al. (2016) [1].

###### S1.1.2 Initialization.

In our implementation, component means are initialized with weighted $k$-means++ (using $\{s_i\}$ as sample weights), and covariances are initialized as scaled identity matrices. Covariance regularization is applied through `reg_covar` in `locat.py`, with default scale

$$\lambda = \max\left(\frac{\text{min\_dist}^{\,2/d}}{6},\, 10^{-4}\right),$$

where min_dist is the mean nearest-neighbor distance in the embedding and $d$ is embedding dimensionality.

Multiple initializations are run in parallel, and the solution with the largest weighted log-likelihood is retained.

###### S1.1.3 Component selection via BIC.

To determine mixture complexity (hyperparameter $K$), we use the Bayesian Information Criterion (BIC), which balances fit quality against model complexity through a parameter-count penalty. In `locat.py`, BIC search is run at two waypoints: full sample size and $\sqrt{n}$ sample size. Candidate component counts start at $K = 1$, and the search range is expanded when the minimum occurs near the current upper boundary. The selected waypoint values are then interpolated to estimate component count as a smooth function of sample size.

###### S1.1.4 Heuristic selection of per-gene component count

In the Locat workflow, a `LOCAT` object is instantiated with expression data and embedding coordinates. For each dataset, a background WGMM over all cells is fit once and reused across gene-wise scanning (e.g., `gmm_scan`). This model partitions the embedding into $K_0$ mixture components, each representing a manifold region. For each gene, we then select a gene-specific component count $K_g$ based on how that gene's weighted cells are distributed across the background components.

Specifically, for each background component we compute the fraction of the gene's total weight assigned to that component (via background responsibilities). We sort these fractions in decreasing order and choose the smallest number of components whose cumulative fraction exceeds a fixed threshold (e.g., 95%). That number defines $K_g$ for the gene's WGMM. Genes concentrated in a few compact regions receive small $K_g$, whereas genes spread across multiple regions receive larger $K_g$. The raw count is then dampened to $\lfloor \sqrt{n+1} \rfloor + 1$, a value that grows much more slowly than the raw count as $n$ increases. This dampened value is clamped to lie between 1 and the smaller of (a) the background GMM's total number of components and (b) one component per 5 assigned cells. The clamp ensures the number of GMM components $K_g$ never exceeds what the background model or the available data can support.

#### S1.2 Empirical null distribution calibration as a function of sample size.

Given a raw concentration statistic $z$, we calibrate it with a Monte Carlo null that is matched to gene sample size.

We define a grid over the fraction of non-zero entries in the sample weight vector:

$$p \in \{p_1, \ldots, p_L\} \subset \big[10/n,\, 1\big], \tag{7}$$

where $n$ is the total number of data points and $L$ is the number of grid points.

Then, for each value of $p$, we repeat the following steps many times:

1. Randomly generate a binary weight vector $\hat{s} \in \{0,1\}^n$ with $np$ non-zero entries.

2. Fit a WGMM using exactly the same pipeline used for real genes.

3. Compute the same raw statistic $z$.

This gives a null distribution of $z$ at each $p$. From that distribution, we estimate the null mean $\mu(p_\ell)$ and null standard deviation $\sigma(p_\ell)$. To evaluate these quantities at arbitrary sample sizes, we fit piecewise cubic Hermite interpolating polynomials (PCHIP) to $\{\mu(p_\ell)\}$ and $\{\sigma(p_\ell)\}$, yielding continuous functions $\mu(p)$ and $\sigma(p)$.

For a real gene with observed proportion $p$ of non-zero weights and raw statistic $z$, we report the calibrated score

$$Z_{\text{null}}(p; z) \;=\; \frac{z - \mu(p)}{\sigma(p)}, \tag{8}$$

which expresses $z$ in standard-deviation units relative to the null at that sample size.

#### S1.3 Global correction and empirical calibration

In Locat, per-gene localization tests are not an exchangeable family of independent tests. Genes differ in prevalence, detection noise, and effective sample size, and each final score combines concentration and depletion components with distinct null behavior. Gene-level statistics are also coupled through shared embedding and model fitting, so the pipeline is not equivalent to i.i.d. draws from one null family. Therefore, we do not use a single genome-wide correction as the primary calibration strategy. Instead, we assess calibration empirically using simulation-based null analyses matched to the pipeline structure and prevalence regime (Supplementary Fig. S1).

#### S1.4 Stability of the Adjusted Localization Score

The adjusted localization score (Eq. 20) includes two penalty coefficients, $\alpha_{\text{size}}$ and $\alpha_{\text{sens}}$, modulating the prevalence and sensitivity components, respectively. Across all datasets we used fixed values $\alpha_{\text{size}} = 0.05$ and $\alpha_{\text{sens}} = 0.12$, selected once based on preliminary simulations and not tuned per dataset.

##### S1.4.1 Sensitivity penalty ($\alpha_{\text{sens}}$).

Fig. S22(A–B) illustrates the effect of $\alpha_{\text{sens}}$ using simulated genes with increasing bleed fraction (fraction of expressing cells outside the primary enriched region). Panel S22A shows representative simulated patterns across the bleed range, and Panel S22B summarizes the resulting $-\log_{10}$ adjusted $p$-values across simulations.

In structured embeddings, many genes show some autocorrelation relative to a weak or random null, so the concentration test can assign similarly small $p$-values to many genes. $\alpha_{\text{sens}}$ refines ranking in this regime by downweighting genes with expression outside enriched regions, thereby reordering genes by bleed. Rankings remain highly stable across $\alpha_{\text{sens}} \in [0.05, 0.30]$ (Spearman $\rho > 0.9998$).

##### S1.4.2 Prevalence penalty ($\alpha_{\text{size}}$).

Supplementary Fig. S22(C–D) shows the effect of $\alpha_{\text{size}}$ across simulated genes spanning a range of prevalences and signal strengths. Panel S22C shows representative simulated patterns across the prevalence range, and Panel S22D summarizes the resulting $-\log_{10}$ adjusted $p$-values.

Increasing $\alpha_{\text{size}}$ lowers adjusted scores across genes, with a larger reduction for low-prevalence genes than for higher-prevalence genes (Panel S22D). This term primarily separates genes with comparable enrichment strength but differing numbers of expressing cells, and prevents very rare genes from filtering to the top when many genes exhibit similarly strong concentration signal. Rankings remain stable across $\alpha_{\text{size}} \in [0.05, 0.20]$, a range spanning the default value used throughout this study ($\alpha_{\text{size}} = 0.05$; Spearman $\rho > 0.96$ relative to this default).

#### S1.5 Operational analysis defaults

To ensure reproducibility and avoid ambiguity between implementation-level function defaults and manuscript analysis settings, Table S1 lists the exact parameter values used in the primary Locat analysis path (`gmm_scan`) unless otherwise stated.

Unless explicitly overridden, background density estimation uses uniform per-cell weights in the primary analysis.

#### S1.6 Robustness to Embedding Choice and Random Seed

To assess robustness of Locat v0.1.5 in the IFN-$\beta$-stimulated PBMC dataset, we split stimulated cells into two disjoint one-third subsets and reran preprocessing independently within each subset, including PCA recomputation. We then ran Locat on embeddings defined by 4 PCs, 8 PCs, 12 PCs, and a BiPCA-based setting in which the effective rank was estimated separately within each fold, that number of PCs was used to construct the neighbor graph, and a UMAP embedding was then computed for Locat. For each fixed embedding, Locat was rerun with three random seeds so that seed variability could be separated from split-to-split variability. Tables S3–S5 summarize cross-fold reproducibility, within-fold seed stability, and embedding sensitivity for these analyses.

#### S1.7 Computational complexity and runtime profiling

For the current implementation in `locat.py`, runtime is naturally split into two phases:

$$T_{\text{total}}(n, G) \;=\; T_{\text{setup}}(n) + T_{\text{scan}}(n, G). \tag{9}$$

**Setup phase.** `T_setup` includes background-model preparation and concentration-null calibration. In the current code path, setup may access full pairwise cell distances during regularization and component-support heuristics. As a result, setup scaling with cell count is superlinear and can include a quadratic term in $n$. For fixed embedding dimension and fixed runtime hyperparameters, setup is therefore primarily cell-count dominated.

**Scan phase.** `T_scan` runs `gmm_scan` across genes. For fixed embedding dimension and bounded mixture complexity, scan runtime grows approximately linearly with the number of tested genes $G$. It also increases with $n$ because per-gene density evaluation and model fitting become more expensive.

**Runtime profiling protocol.** To make implementation-level scaling explicit and reproducible, we benchmarked runtime on real data from the IFN-$\beta$-stimulated PBMC dataset by subsampling cells and genes over predefined grids.

1. Load one real dataset and fixed embedding coordinates.
2. For each $(n, G)$ pair, subsample $n$ cells and $G$ genes without replacement.
3. Measure setup time separately (background fit + null calibration).
4. Measure scan time separately (`gmm_scan` over selected genes).
5. Repeat each $(n, G)$ condition across multiple random seeds and report median wall-clock time.

Benchmarking was run on a system using an AMD EPYC 7282 16-Core CPU and NVIDIA RTX A6000 GPU.

Empirical scaling is shown in Fig. S23. Panel A (runtime versus number of genes at fixed $n$) shows near-linear growth of scan time with $G$, while setup time is comparatively stable across $G$ at fixed $n$. Panel B (runtime versus number of cells at fixed $G$) shows that setup increases steeply with $n$, and scan also increases with $n$ across all gene-grid settings.

#### S1.8 SpectralRH

##### S1.8.1 Design gap in existing methods

SpectralRH was designed to address a specific methodological gap in existing gene localization methods. We believe that our analyses of the advantages of our dual concentration-depletion criterion would not be complete without addressing this gap.

Existing gene localization methods detect spatial patterns through distinct but complementary approaches.

- Hotspot quantifies local spatial autocorrelation via Geary's C, identifying genes with correlated expression among neighboring cells.
- LMD measures diffusion rates on the cell graph, scoring genes whose expression is concentrated in compact regions where random walks remain localized.
- singleCellHaystack compares kernel density estimates in low-dimensional spatial coordinates, detecting genes enriched in specific physical locations.
- Scanpy's rank_genes_groups applies Wilcoxon rank-sum tests to identify differentially expressed genes between predefined cell groups.
- GiniClust uses Gini index statistics to detect rare cell types through expression dispersion patterns.
- GSPA learns gene embeddings by decomposing expression signals using diffusion wavelets at multiple scales, enabling pattern-based gene clustering across the manifold.

While these methods successfully identify genes marking discrete clusters or spatially enriched regions, they do not distinguish whether localized expression patterns align with the embedding's global geometric structure, e.g., developmental trajectories or continuous cell-state transitions, or instead represent arbitrary spatial subpopulations. This constitutes a substantial methodological gap, particularly because embeddings constructed from transcriptional profiling are often low-dimensional representations of latent cell-state structure and developmental trajectories.

SpectralRH addresses this gap by quantifying expression smoothness and spectral energy concentration in low-frequency Laplacian eigenmodes, explicitly testing whether gene expression patterns follow large-scale manifold geometry. Genes that vary smoothly along trajectories or gradients exhibit high energy in low frequencies, whereas genes marking disconnected clusters or arbitrary regions on the embedding do not, enabling SpectralRH to prioritize trajectory-associated markers over arbitrary localized patterns.

##### S1.8.2 Algorithm

**SpectralRH** quantifies how smoothly and coherently each gene's expression varies across the cell-state manifold. Let $X \in \mathbb{R}^{n \times G}$ be a normalized single-cell expression matrix, where $n$ is the number of cells and $G$ is the number of genes. Let $L = I - D^{-1/2} W D^{-1/2}$ be the symmetric normalized Laplacian of the cell-cell connectivity graph (constructed from a $k$-nearest-neighbor graph in PCA space). Each gene expression profile $x_g \in \mathbb{R}^n$ is treated as a graph signal.

For each gene $g$, we compute two complementary spectral quantities:

1. **Rayleigh smoothness:**
$$R_g = \frac{x_g^\top L x_g}{x_g^\top x_g}, \tag{10}$$
which measures the graph-smoothness of $x_g$—i.e., how rapidly expression changes between neighboring cells. Smaller values of $R_g$ correspond to smoother, spatially coherent signals.

2. **Spectral entropy:**
$$H_g = -\sum_{k=1}^{m} p_{gk} \log p_{gk}, \quad p_{gk} = \frac{|\hat{x}_{gk}|^2}{\sum_{\ell=1}^{m} |\hat{x}_{g\ell}|^2}, \tag{11}$$
where $\hat{x}_g = U^\top x_g$ are the coefficients of $x_g$ in the first $m$ eigenvectors $U$ of $L$ (corresponding to the smallest eigenvalues). $H_g$ measures how concentrated the signal's energy is in the low-frequency eigenmodes of the graph; low-entropy signals have compact, structured spatial support.

Because both $R_g$ and $H_g$ depend strongly on the number of expressing cells, we constructed sample-size-matched null distributions. For each observed expression prevalence $k_g = \sum_i \mathbb{I}[x_{ig} > 0]$, we generated random expression patterns with the same sparsity and computed corresponding null values $(R_g^0, H_g^0)$. Observed scores were then converted to empirical $p$-values or percentiles relative to these nulls, improving comparability across genes with different prevalence.

Finally, for each gene we normalized and averaged the two calibrated metrics to produce a single composite score:
$$\text{SpectralRH}_g = \tfrac{1}{2}\left(\tilde{R}_g + \tilde{H}_g\right), \tag{12}$$
where $\tilde{R}_g$ and $\tilde{H}_g$ denote normalized, size-matched percentiles of Rayleigh smoothness and spectral entropy, respectively. Lower values of the SpectralRH score indicate smoother and more structured expression patterns consistent with spatial localization.

#### S2 Supplementary Data

CSV files used to populate the split-stability summary tables for the IFN-$\beta$ stimulation analysis, together with the full ESC time-course localization results table, are included as:

- `data/stim_twothirds_cv_table_across_fold_wide.csv`
- `data/stim_twothirds_cv_table_within_seed_wide.csv`
- `data/stim_twothirds_cv_table_pc_sensitivity_wide.csv`
- `data/esc_locpatterns_dataframe.csv`

### S3 Supplementary Figures

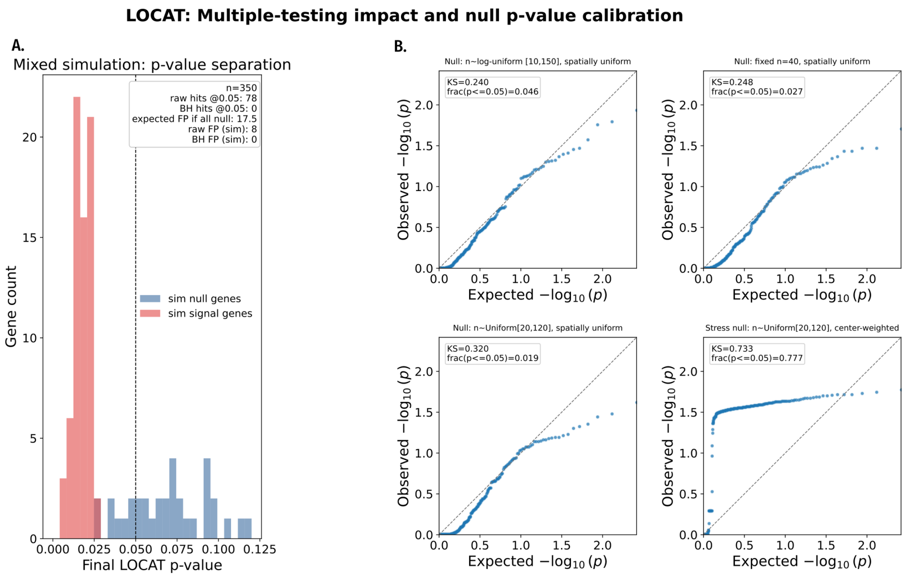

Figure S1: Simulation-based calibration of Locat localization $p$-values in a synthetic 2D Gaussian embedding (4500 cells). (**A**) Mixed simulation with 350 genes (70 signal, 280 null; prevalence 25–89 cells per gene). Signal genes are enriched near the embedding center and null genes are sampled spatially uniformly. Localization $p$-values separate signal from null genes, while BH correction is overly conservative. (**B**) QQ plots under four null-only settings (260 genes each): fixed $n = 40$, $n \sim$ Uniform[20120], log-uniform on [10150], and a null with expression biased toward the embedding center. Uniform scenarios track nominal calibration; the spatially biased null shows the largest deviation, as null genes are concentrated in the same region as signal genes.

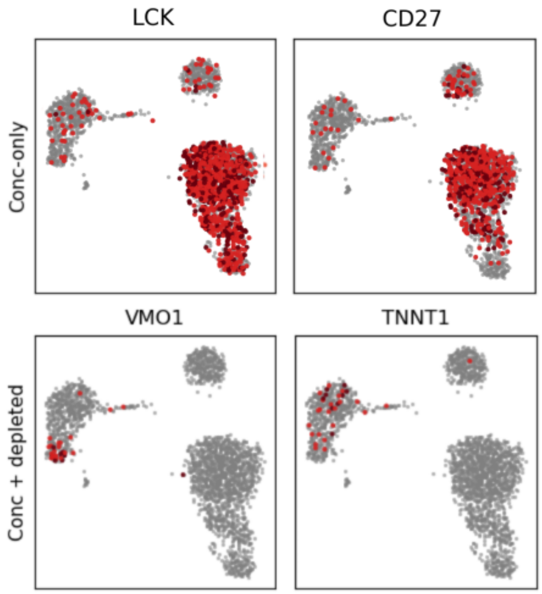

Figure S2: UMAP embeddings for the Scanpy PBMC3k dataset, grouped into concentration-only genes (top) and genes classified as significantly localized by combined concentration and depletion criteria (bottom). Genes in the bottom row show minimal expression outside enriched regions while varying in regional density and compactness.

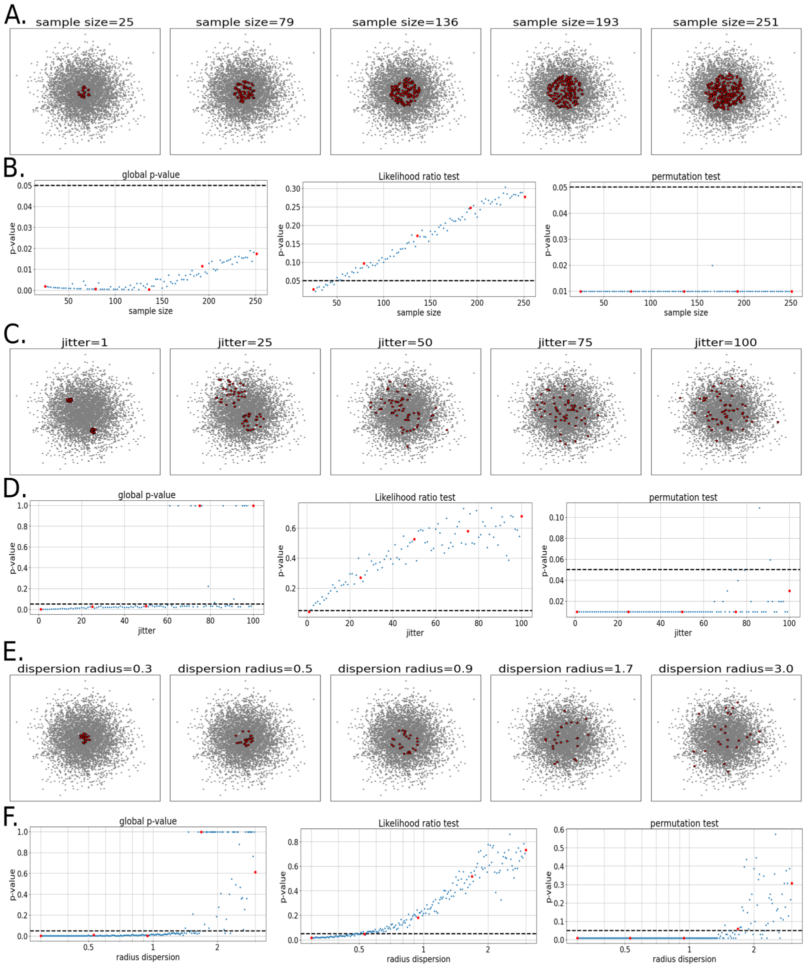

Figure S3: Validation of Locat on simulated data with controlled ground truth. (A, C, E) UMAP embeddings of 1500 cells from a Gaussian background (gray points, contour lines) with five replicates of a simulated localized gene under three conditions: (A) sample size (25–251 expressing cells), (C) positional jitter (1–100% of the maximum tested jitter), and (E) dispersion radius (0.3–3.0 units). (B, D, F) Scatter plots of $p$-values from 50 replicates per condition for global Locat (left), likelihood ratio test (middle), and permutation-based Locat (right). Dashed lines indicate $\alpha = 0.05$.

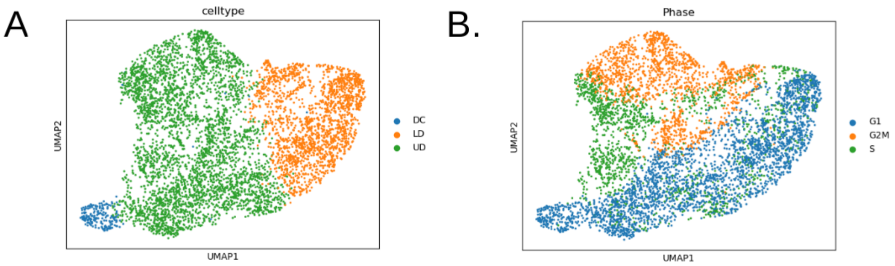

Figure S4: Murine embryonic dermis dataset at E14.5, containing approximately 5000 cells spanning the transition from upper dermal fibroblasts to Sox2-positive dermal condensates (early hair-follicle precursors). This system forms a continuous transcriptional manifold without discrete cluster boundaries. **(A, B)** t-SNE embedding constructed from 4000 highly variable genes and labeled by dermal condensate (DC) versus upper dermal (UD) cell type (**A**) and by primary cell-cycle phase (**B**).

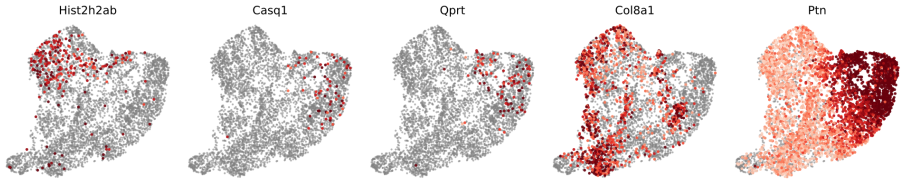

Figure S5: Representative genes with large rank discrepancies between Locat and other methods. Genes ranked highly by Locat but lower by other methods (*Hist1h2bb*, *Casq1*, *Qprt*) show tightly confined expression with minimal background (e.g., *Casq1*: Locat 99th percentile, Hotspot 64th percentile). Genes ranked higher by enrichment-focused methods (*Col8a1*, *Ptn*) show moderate local enrichment with substantial background expression (e.g., *Ptn*: LMD 99th percentile, Locat 5th percentile). These diffuse patterns score highly under concentration-only criteria but are deprioritized by the depletion term in Locat.

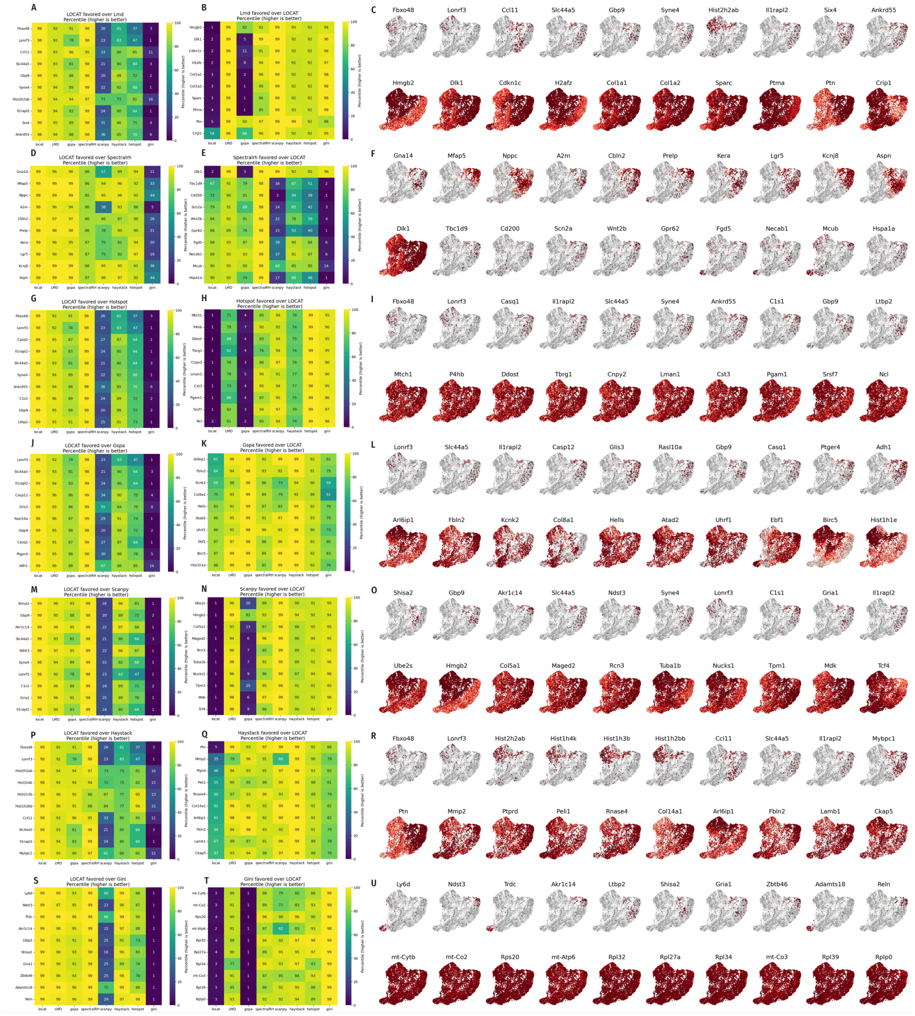

Figure S6: Comparison of Locat with alternative methods using the same gene set and preprocessed murine developmental dermis data. Percentile discrepancies are summarized for each method pair. Panels **A**–**C** show the Locat versus LMD comparison: genes favored by Locat (**A**), genes favored by LMD (**B**), and expression plots for those genes (**C**). The same panel structure is repeated for SpectralRH (**D**–**F**), Hotspot (**G**–**I**), GSPA (**J**–**L**), Scanpy (**M**–**O**), singleCell-Haystack (**P**–**R**), and Gini (**S**–**U**).

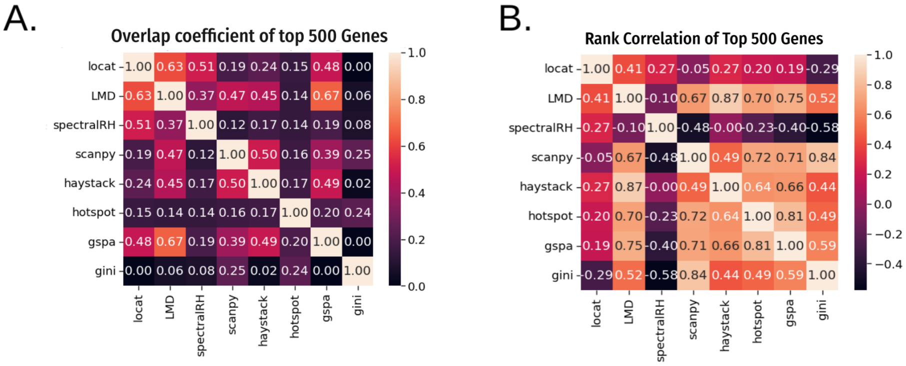

Figure S7: Restricting analysis to the top 500 genes per method still shows that other rankings remain distinct from Locat. **A.** Overlap coefficient for top-500 gene sets across methods; LMD and SpectralRH show the highest overlap with Locat (0.63 and 0.51, respectively). **B.** Correlation matrix computed on top 500 genes ranked by Locat scores; LMD has the highest correlation again in this regime.

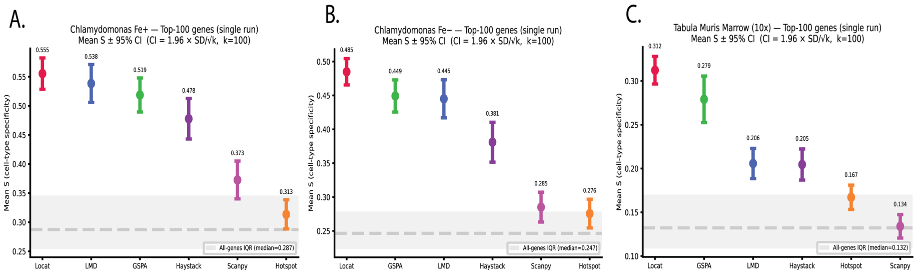

Figure S8: Cell-type specificity of top-100 genes in three additional datasets. Mean cell-type specificity score $S \pm 95\%$ CI $(1.96 \times \mathrm{SD}/\sqrt{k}$, $k = 100)$ for top-100 genes returned by each method. Methods are ordered left to right by decreasing mean $S$ within each panel. Gray shaded region: interquartile range of $S$ across all genes; dashed line: all-genes median $S$. **A.** *Chlamydomonas reinhardtii* iron-sufficient ($Fe^{+}$) transcriptome [2]; cell-type groupings are Leiden clusters (resolution 0.3, 5 clusters). **B.** Same dataset, iron-deficient ($Fe^{-}$) condition (6 clusters). **C.** Murine bone marrow from the Tabula Muris Senis 10x dataset (single donor) [3]. Locat ranks first in mean cell-type specificity in all three datasets, consistent with the main comparison (Fig. 2).

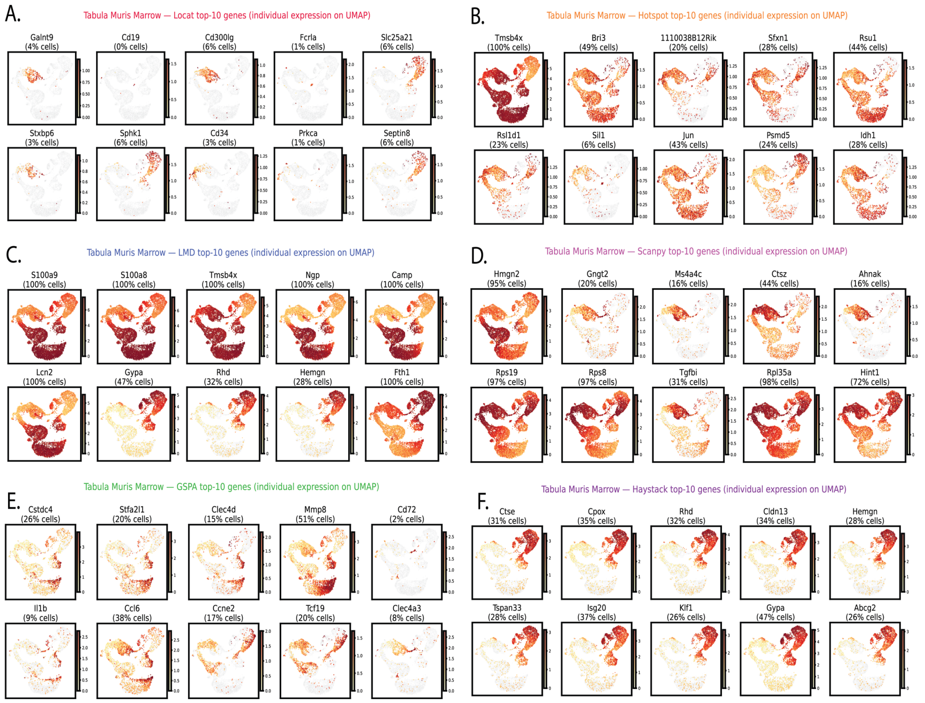

Figure S9: Top-10 gene expression on UMAP for the Tabula Muris Senis bone marrow dataset [3]. Each panel shows individual gene expression for the top-10 genes returned by one method (log-normalized counts; yellow–orange–red scale); non-expressing cells are shown in gray. The percentage of cells with non-zero expression is shown in parentheses. **(A–F)** Top-10 genes for Locat, Hotspot, LMD, Scanpy, GSPA, and singleCellHaystack, respectively. Locat top-10 genes are expressed in 0–6% of cells and show visibly constrained background expression, mapping to spatially restricted regions of the embedding.

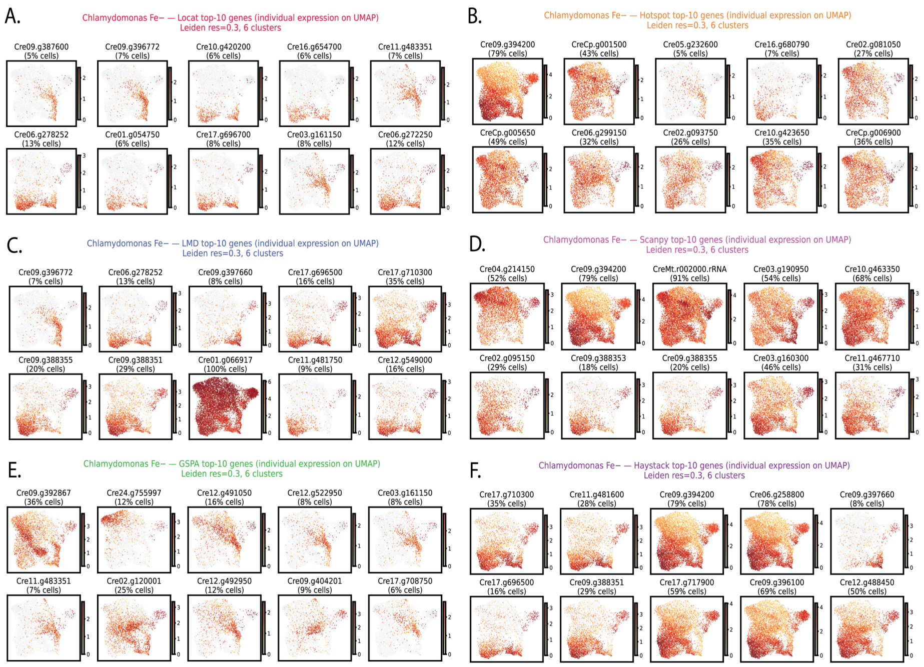

Figure S10: Top-10 gene expression on UMAP for the *Chlamydomonas reinhardtii* iron-deficient ($Fe^{-}$) transcriptome [2]. Each panel shows individual gene expression for the top-10 genes returned by one method (log-normalized counts; yellow–orange–red scale); non-expressing cells are shown in gray. The percentage of cells with non-zero expression is shown in parentheses; Leiden clustering at resolution 0.3 (6 clusters). **(A–F)** Top-10 genes for Locat, Hotspot, LMD, Scanpy, GSPA, and singleCellHaystack, respectively. Locat top-10 genes are expressed in 5–13% of cells with noticeably constrained background expression.

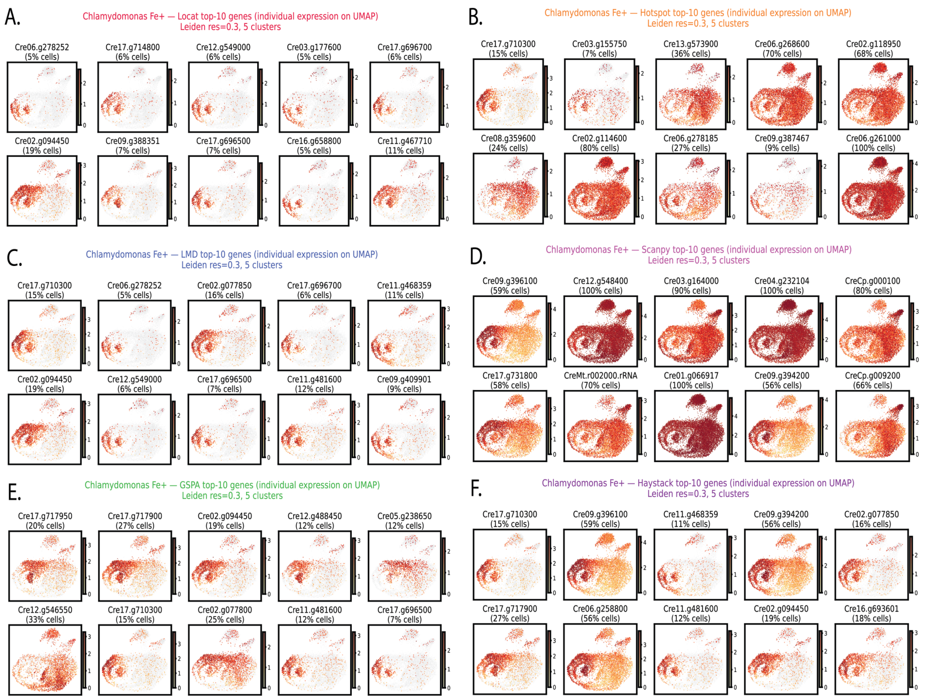

Figure S11: Top-10 gene expression on UMAP for the *Chlamydomonas reinhardtii* iron-sufficient ($Fe^{+}$) transcriptome [2]. Each panel shows individual gene expression for the top-10 genes returned by one method (log-normalized counts; yellow–orange–red scale); non-expressing cells are shown in gray. The percentage of cells with non-zero expression is shown in parentheses; Leiden clustering at resolution 0.3 (5 clusters). **(A–F)** Top-10 genes for Locat, Hotspot, LMD, Scanpy, GSPA, and singleCellHaystack, respectively. Locat top-10 genes are expressed in 5–19% of cells with noticeably constrained background expression.

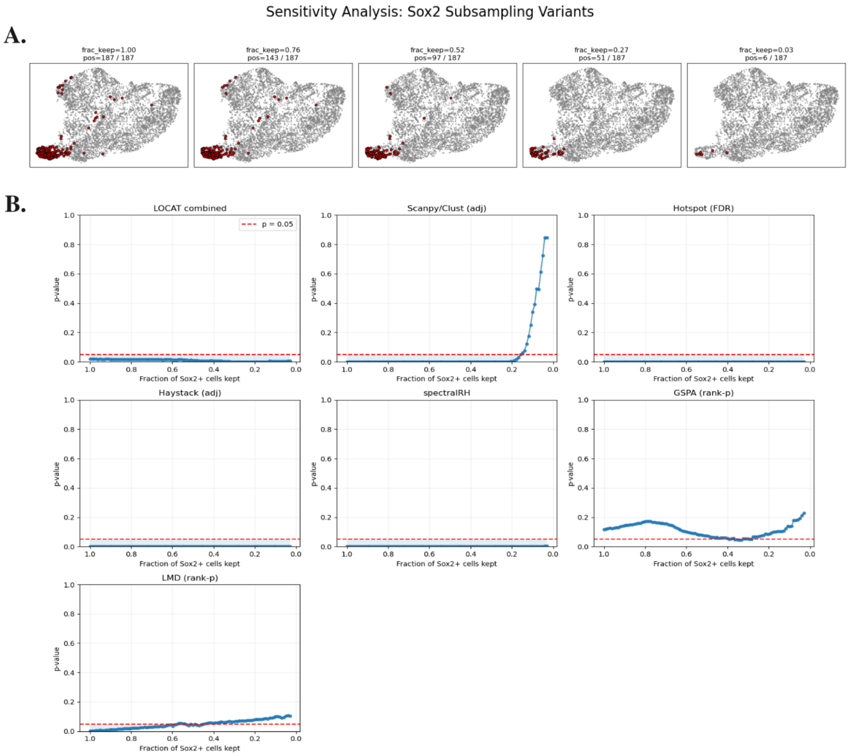

Figure S12: (A) UMAP embeddings of the murine dermis dataset showing simulated *Sox2* expression under progressive subsampling of expressing cells, with the original expression pattern and cell embedding held fixed. (B) Sensitivity of gene prioritization methods as a function of the fraction of *Sox2*-expressing cells retained. Locat maintains detectability across nearly the full range of subsampling levels. SpectralRH, Hotspot, and singleCellHaystack show limited sensitivity to reduced prevalence, whereas cluster-based differential expression rapidly loses sensitivity as expressing cells become rare. GSPA and LMD exhibit a gradual degradation of empirical significance values with decreasing prevalence.

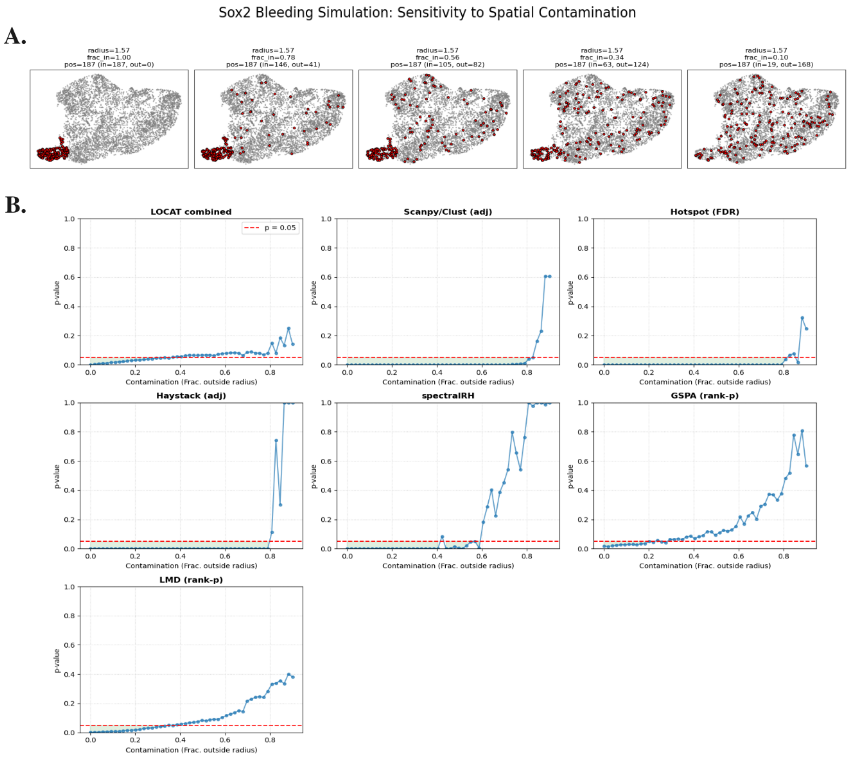

Figure S13: (A) UMAP embeddings showing progressive redistribution of simulated *Sox2* expression into the background population while holding the number of expressing cells fixed. (B) Response of gene prioritization methods to increasing diffusion of *Sox2* expression. Locat shows a graded loss of detectability, with the combined $p$-value increasing and crossing the detection threshold as expression becomes broadly distributed. SpectralRH, GSPA, and LMD similarly exhibit monotonic score increases with increasing diffusion, while Hotspot, singleCellHaystack, and cluster-based differential expression remain largely insensitive until extreme diffusion.

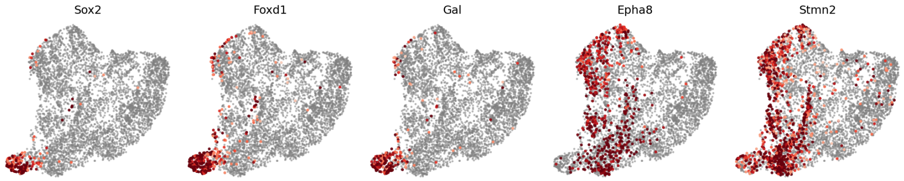

Figure S14: UMAP embeddings colored by expression of representative genes identified as localized by Locat. Dermal condensate-associated genes (*Sox2*, *Foxd1*, *Gal*) localize to terminal regions of the fibroblast-to-dermal condensate trajectory. Cell-cycle-associated genes (*Kif2c*, *Gtse1*) exhibit spatially restricted expression patterns consistent with discrete cell-cycle states.

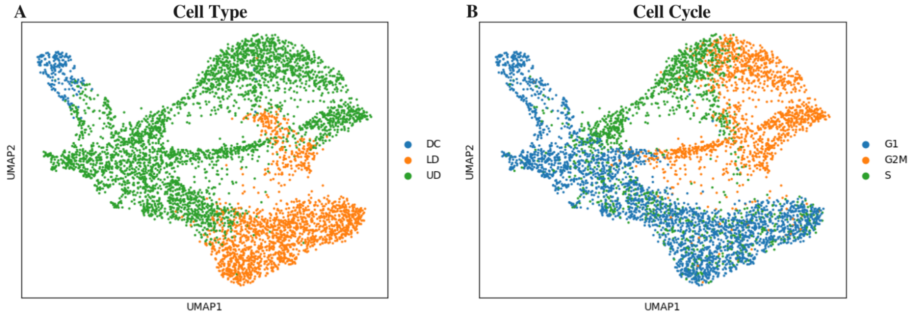

Figure S15: localized-gene embedding (LG-embedding) labeled by (**A**) dermal cell type (UD, LD, DC) and (**B**) cell-cycle phase (G1, G2/M, S). In the murine embryonic dermis dataset, Locat identified 430 genes at $p < 0.05$ with spatially compact expression patterns aligned with proliferative and differentiation-associated variation. This embedding was computed from the significantly localized gene subset.

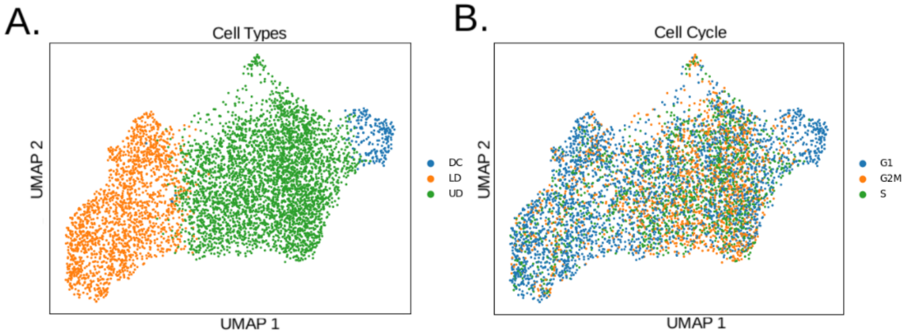

Figure S16: Embedding coordinates computed from a differentiation-focused gene partition in the murine embryonic dermis dataset, as defined in a prior study [4].

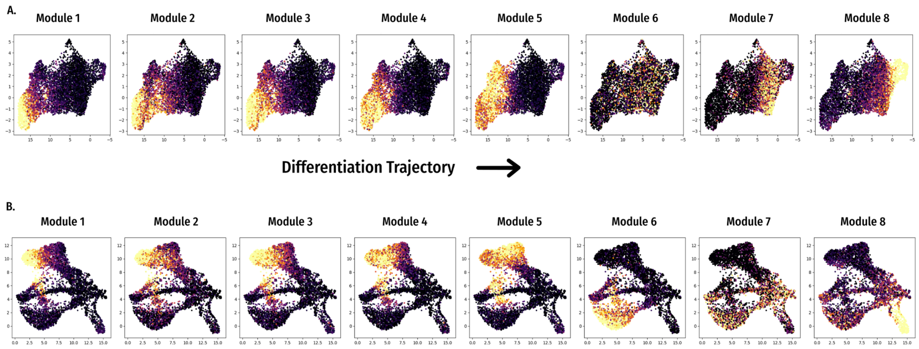

Figure S17: Average module expression for all eight non-singleton localized-gene modules, shown on (**A**) the differentiation-focused UMAP embedding (Fig. S16) and (**B**) the localized-gene embedding (LG-embedding; Fig. S15). Modules were derived from 1302 significantly localized genes ($\alpha = 0.05$) after training Locat on the murine dermal condensate dataset using UMAP coordinates from a differentiation-focused gene partition [4].

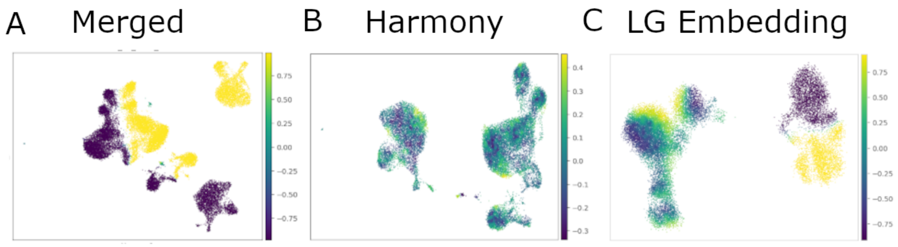

Figure S18: Differential-abundance results projected onto UMAP embeddings generated using (left) a simple merge of preprocessed CTR and STIM samples, (middle) Harmony batch correction, and (right) a simple merge using only localized genes. Batch correction over-smooths the two samples, whereas merging on all genes fully separates them. Using localized genes aligns most shared populations while preserving separation of IFN-responsive populations [5].

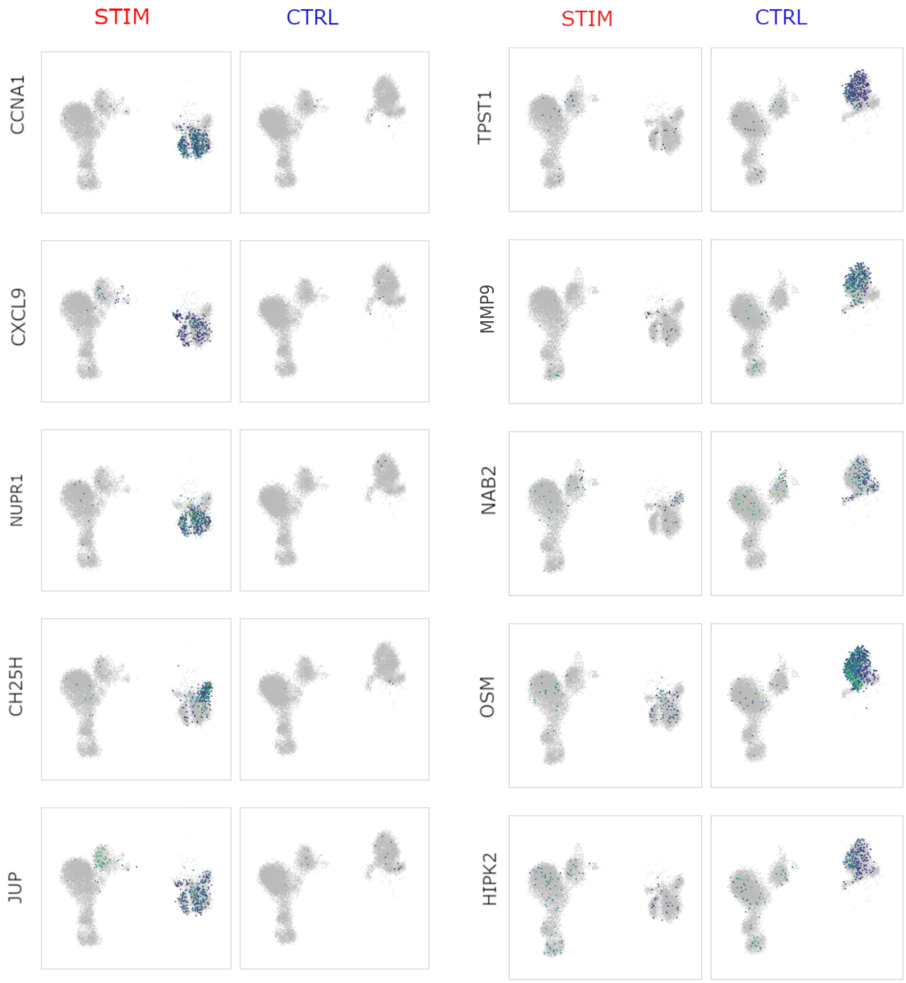

Figure S19: Condition-specific localization examples plotted on a local-gene embedding that aligns CTR and STIM samples from the IFN-$\beta$ stimulation dataset [5].

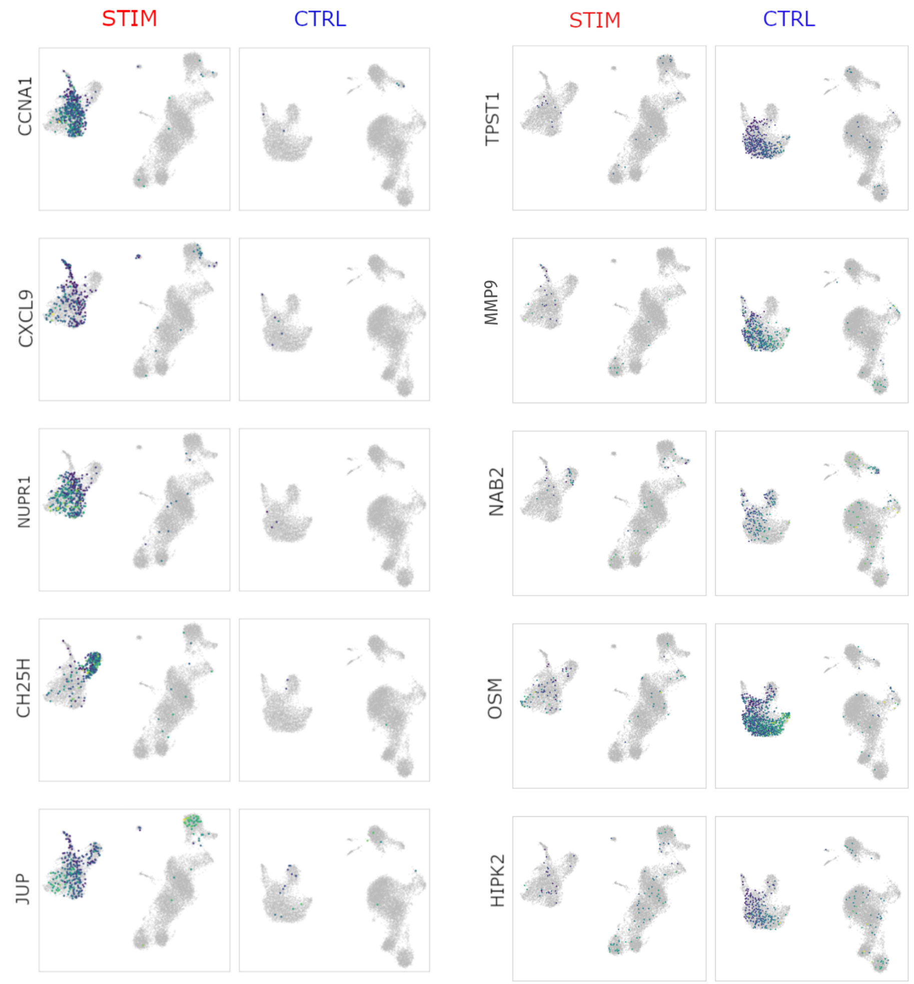

Figure S20: Condition-specific localization examples plotted on per-sample UMAP embeddings from the IFN-$\beta$ stimulation dataset [5].

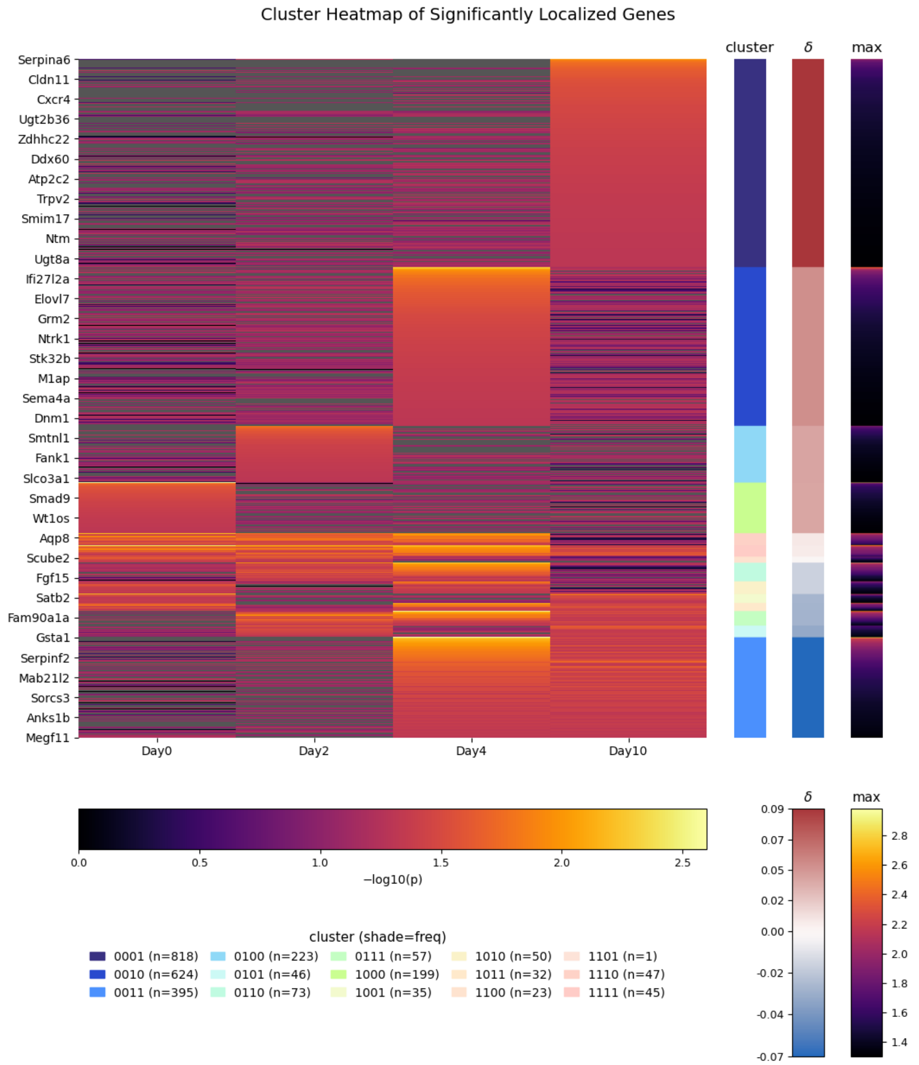

Figure S21: Heatmap of combined localization $p$-values across time points for all genes (rows) that pass a significance threshold in at least one time point. Colors follow the lower color bar; NA values (unobserved genes) are shown in gray. Genes are grouped by shared temporal localization patterns. The right-side bars report each gene's maximum $-\log_{10}$ localization $p$-value (overall significance), $\delta$ (deviation of observed versus expected pattern frequency), and pattern-cluster identity.

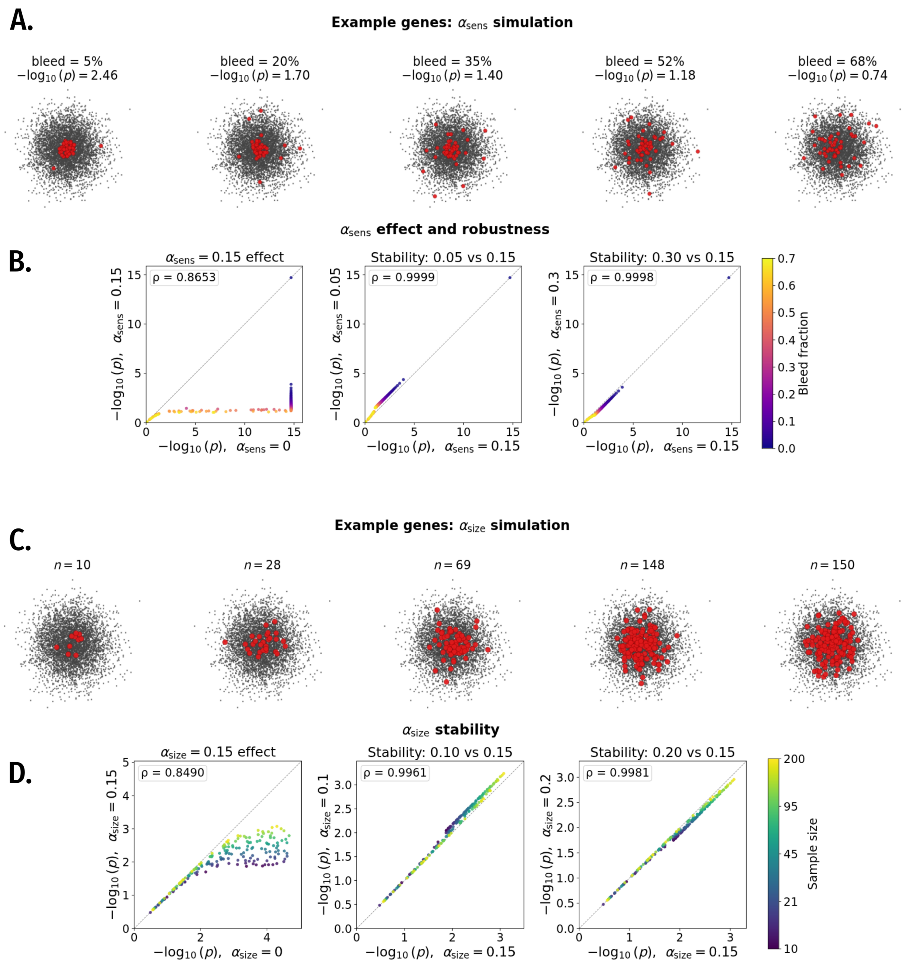

Figure S22: Effect of adjustment coefficients in a 5000-cell Gaussian embedding. (**A**–**B**) Sensitivity coefficient ($\alpha_{\text{sens}}$): genes simulated at fixed prevalence ($n = 50$) with 0–70% bleed outside a focal enriched region. (**A**) Representative simulated patterns and (**B**) corresponding $-\log_{10}$ adjusted $p$-values. Increasing $\alpha_{\text{sens}}$ progressively downweights high-bleed genes while preserving overall ranking stability. (**C**–**D**) Prevalence coefficient ($\alpha_{\text{size}}$): genes simulated across a log-uniform range of expressing cell counts (10–200). Increasing $\alpha_{\text{size}}$ penalizes low-prevalence genes more strongly, separating genes with similar enrichment but differing cell counts. Rankings remain stable across the tested parameter ranges.

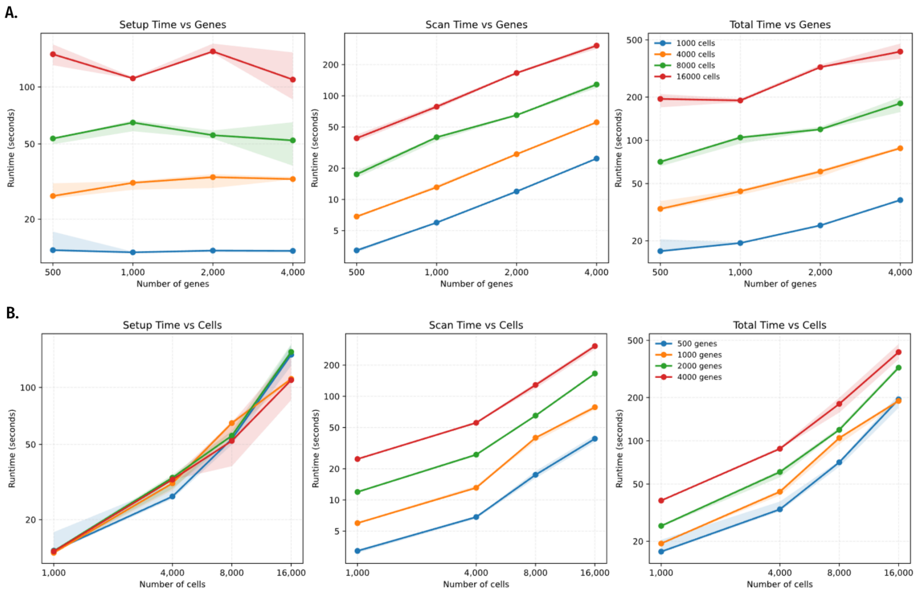

Figure S23: Runtime scaling for Locat using the current `locat.py` implementation. **A.** Setup, scan, and total runtime versus number of genes (fixed cell count). **B.** Setup, scan, and total runtime versus number of cells (fixed gene count). Benchmarks used three replicates per grid point and report median trends with variability bands.

### S4 Supplementary Tables

Table S1: Primary analysis defaults used for Locat (gmm_scan).

| Parameter | Value | Role |
|---|---|---|
| max_freq | 0.9 | Exclude genes above prevalence threshold from concentration/depletion testing. |
| rc_weight_mode | binary | Depletion observed fraction uses expressing-cell support (not expression magnitude). |
| rc_rho_bb | 0.02 | Beta-binomial overdispersion for depletion tail probability. |
| rc_lambda_values | [1.0, 2.0] (8 points, linearly spaced) | Contrast-threshold scan grid for depletion. |
| rc_min_p0_abs | 0.10 | Minimum background mass in depletion region. |
| rc_min_expected | 3 | Minimum expected effective count in depletion region. |
| rc_min_abs_deficit | 0.04 | Minimum absolute depletion deficit. |
| rc_eps_rel | 0.01 | Relative contrast guard in depletion scan. |
| rc_n_eff_scale | 0.6 | Multiplicative scaling applied to Kish effective sample size. |
| rc_n_trials_cap | $\sqrt{n_{\text{cells}}}$ (rounded to integer) | Upper cap on effective trials for depletion test. |
| rc_p_floor | $10^{-12}$ | Numerical floor for depletion $p$-values. |
| rc_soft_bound | 1.0 | Optional upper bound on depletion-region background mass ($p0$_abs); active only when set below 1.0. |
| $\alpha_{\text{size}}$ | 0.05 | Sparsity penalty coefficient in adjusted localization score. |
| $\alpha_{\text{sens}}$ | 0.12 | Sensitivity penalty coefficient in adjusted localization score. |

Table S2: Difference between observed and expected frequencies ($\delta$) for each temporal localization cluster in the time series of embryonic stem cell differentiation following retinoic acid induction. The four digits of each cluster label correspond to the four time points, with the first digit corresponding to Day 0, the second to Day 2, and so on. Each digit is binary, with values of 1 in a given label and digit indicating localization is detected for that cluster at that time point.

| cluster label | $\delta$ |
|---|---|
| 0001 | 0.090 |
| 0010 | 0.049 |
| 0100 | 0.039 |
| 1000 | 0.038 |
| 1110 | 0.009 |
| 1111 | 0.007 |
| 1100 | -0.000 |
| 1101 | -0.010 |
| 0110 | -0.017 |
| 1010 | -0.017 |
| 1001 | -0.028 |
| 1011 | -0.029 |
| 0111 | -0.030 |
| 0101 | -0.035 |
| 0011 | -0.065 |

Table S3: Cross-fold reproducibility of Locat gene localization rankings in the IFN-$\beta$ stimulation dataset. Two disjoint one-third subsets of stimulated cells were analyzed independently, with PCA recomputed within each fold. For the BiPCA setting, effective rank was estimated per fold (47 for fold A, 48 for fold B) and used to construct the UMAP embedding. The AUROC row reports recovery of significant genes via held-out $-\log_{10}(p + 10^{-12})$ scores across the shared gene panel. Entries report means with 95% CI half-widths.

| **Metric** | **4 PCs** | **8 PCs** | **12 PCs** | **BiPCA→UMAP** |
|---|---|---|---|---|
| AUROC (all genes; $-\log_{10} p$) | 0.923 ± 0.009 | 0.977 ± 0.004 | 0.966 ± 0.004 | 0.979 ± 0.003 |
| Spearman rho (top 200) | 0.862 ± 0.104 | 0.807 ± 0.081 | 0.728 ± 0.065 | 0.824 ± 0.064 |
| Spearman rho (top 500) | 0.697 ± 0.087 | 0.762 ± 0.034 | 0.681 ± 0.009 | 0.809 ± 0.028 |
| Jaccard overlap (significant genes) | 0.694 ± 0.043 | 0.740 ± 0.030 | 0.731 ± 0.063 | 0.734 ± 0.031 |

Table S4: Within-fold seed stability of Locat gene localization rankings in the IFN-$\beta$ stimulation dataset. For each fold and embedding setting, Locat was rerun with three random seeds with the embedding held fixed. Entries summarize all within-fold seed-pair comparisons across the two folds. For the BiPCA setting, effective rank was estimated per fold (47 for fold A, 48 for fold B) and used to construct the UMAP embedding. The AUROC row reports recovery of significant genes via held-out $-\log_{10}(p + 10^{-12})$ scores. Values report means with 95% CI half-widths.

| **Metric** | **4 PCs** | **8 PCs** | **12 PCs** | **BiPCA→UMAP** |
|---|---|---|---|---|
| AUROC (all genes; $-\log_{10} p$) | 0.940 ± 0.013 | 0.978 ± 0.005 | 0.969 ± 0.009 | 0.985 ± 0.007 |
| Spearman rho (top 200) | 0.876 ± 0.036 | 0.832 ± 0.047 | 0.754 ± 0.065 | 0.921 ± 0.024 |
| Spearman rho (top 500) | 0.748 ± 0.053 | 0.792 ± 0.012 | 0.709 ± 0.025 | 0.874 ± 0.046 |
| Jaccard overlap (significant genes) | 0.773 ± 0.036 | 0.743 ± 0.026 | 0.769 ± 0.018 | 0.829 ± 0.040 |

Table S5: Embedding sensitivity of Locat gene localization rankings in the IFN-$\beta$ stimulation dataset. Entries summarize within-fold comparisons between embedding settings, pooled across both folds and all seed-matched pairs. For the BiPCA setting, effective rank was estimated per fold (47 for fold A, 48 for fold B) and used to construct the UMAP embedding. The AUROC row reports recovery of significant genes via held-out $-\log_{10}(p+10^{-12})$ scores. Values report means with 95% CI half-widths.

| **Metric** | **4 vs 8** | **4 vs 12** | **4 vs BiPCA** | **12 vs BiPCA** |
|---|---|---|---|---|
| AUROC (all genes; $-\log_{10} p$) | 0.934 ± 0.014 | 0.921 ± 0.030 | 0.933 ± 0.016 | 0.951 ± 0.024 |
| Spearman rho (top 200) | 0.778 ± 0.035 | 0.684 ± 0.057 | 0.758 ± 0.047 | 0.748 ± 0.040 |
| Spearman rho (top 500) | 0.673 ± 0.026 | 0.592 ± 0.031 | 0.685 ± 0.073 | 0.682 ± 0.040 |
| Jaccard overlap (significant genes) | 0.526 ± 0.029 | 0.435 ± 0.046 | 0.561 ± 0.035 | 0.624 ± 0.055 |

### S5 Supplementary Notes
